# Elevated p21 (CDKN1a) mediates β-thalassemia erythroid apoptosis but its loss does not improve β-thalassemic erythropoiesis

**DOI:** 10.1101/2022.03.03.482874

**Authors:** Vijay Menon, Miao Lin, Raymond Liang, Tasleem Arif, Anagha Menon, Laura Breda, Stefano Rivella, Saghi Ghaffari

**Author notes:** **Correspondence:** Saghi Ghaffari, M.D.-Ph.D., Department of Cell, Developmental & Regenerative Biology, Icahn School of Medicine at Mount Sinai, New York, NY 10029. Current position: Department of Therapeutic Radiology, Yale University, Connecticut (Vijay Menon). Current position: HemoGenyx Pharmaceuticals (Raymond Liang). These authors contributed equally to the work (Vijay Menon, Miao Lin, Raymond Liang).

## Abstract

β-thalassemias are common hemoglobinopathies due to mutations in the β-globin gene that lead to hemolytic anemias. Premature death of β-thalassemic erythroid precursors results in ineffective erythroid maturation, increased production of erythropoietin (Epo), expansion of erythroid progenitor compartment, extramedullary erythropoiesis and splenomegaly. However, the molecular mechanism of erythroid apoptosis in β-thalassemia is not well understood. Using a mouse model of β-thalassemia (*Hbb*^*th3/+*^), we show that dysregulated expression of Foxo3 transcription factor and its upstream pro-apoptotic regulator TP53 is implicated in β-thalassemia erythroid apoptosis. In *Foxo3*^*-/-*^ */Hbb*^*th3/+*^ mice, erythroid apoptosis is significantly reduced while erythroid cell maturation, red blood cell and hemoglobin production are substantially improved. However, persistence of elevated reticulocytes and splenomegaly suggests that ineffective erythropoiesis is not resolved in *Foxo3*^*-/-*^*/Hbb*^*th3/+*^. We next focused on cell cycle inhibitor Cdkn1a (p21) and show that p21 that is a target of both Foxo3 and TP53 is markedly upregulated in both mouse and patients-derived β-thalassemic erythroid precursors. To address the contribution of p21 to β-thalassemia pathophysiology, we generated *p21*^*-/-*^ */Hbb*^*th3/+*^ mice. Double mutant p21*/Hbb*^*th3/+*^ mice exhibited embryonic lethality with only a fraction of mice surviving to weaning. Notably, studies in adult mice showed apoptosis and circulating Epo were greatly reduced in erythroid compartments of surviving *p21*^*-/-*^ */Hbb*^*th3/+*^ relative to *Hbb*^*th3/+*^ mice, while ineffective erythroid cell maturation, extramedullary erythropoiesis and splenomegaly were not modified. These combined results indicate that while lack of Foxo3 reduces apoptosis and improves anemia, diminished p21-mediated apoptosis is insufficient to improve red blood cell production in *Hbb*^*th3/+*^ mice. They also suggest that a molecular network constituted by p21, FOXO3 and TP53, control erythroid cell survival and differentiation in β-thalassemia. Overall, these studies provide a new framework for investigating ineffective erythropoiesis in β-thalassemia.

**Key Points:**

- Elevated p21 mediates β-thalassemia erythroid cell apoptosis
- Loss of Foxo3 or p21 reduces β-thalassemia erythroid cell apoptosis but does not improve β-thalassemia ineffective erythropoiesis
- A network of Foxo3, p21 and TP53 controls β-thalassemia erythroid apoptosis
- Apoptosis may be uncoupled from ineffective erythropoiesis in β-thalassemia

## Introduction

β-thalassemias are one of the most common hemoglobinopathies in the world caused by over 200 mutations in the β-globin gene. Patients suffer from variable degrees of anemia associated with a broad range of clinical manifestations ^1-3^. Patients with ß-thalassemia may be affected by severe anemia, requiring chronic blood transfusions for survival (β-thalassemia major), or milder clinical anemia requiring only sporadic transfusions (β-thalassemia intermedia), while patients with minor thalassemia are asymptomatic ^1-3^.

Mutations in β-globin gene lead to the absence or reduced β globin synthesis causing an imbalance in the α vs. β chain – of the adult α_2_β_2_ hemoglobin A (HbA) – that is mostly evident in homozygote β-thalassemia mutations. The accumulation of excessive unpaired α-globin chains in erythroblasts triggers redox-mediated reactions leading to the generation of toxic aggregates that precipitate into red blood cells (RBCs) precursors, and result in premature death, ineffective erythroid maturation and hemolytic anemia. The β-thalassemia anemia triggers the release of erythropoietin (Epo) which leads to the expansion of greatly Epo-sensitive erythroid progenitors, extramedullary erythropoiesis and splenomegaly. The increased apoptosis of late maturing erythroblasts is known as an aggravating factor in sustaining anemia despite enhanced erythropoiesis in β-thalassemic patients ^4-6^, which results in further expansion of extramedullary hematopoiesis. The clinical aspects of β-thalassemia are orchestrated in response to stress and elevated reactive oxygen species (ROS) in erythroid cells ^7-11^. However, the source of elevated ROS levels remains debated and whether ROS are implicated in regulating erythroid apoptosis in β-thalassemia is unclear ^12^. Importantly, the molecular underpinning of the enhanced apoptosis in β-thalassemia remains poorly understood. ^4-6,12-14^.

In erythroid compartment, the transcription factor Foxo3 coordinates cell maturation and redox state ^15-19^. Foxo3 is required for homeostatic erythropoiesis and optimum gene expression during terminal erythroid maturation and enucleation ^14,15^. These diverse functions are accomplished through Foxo3’s direct transcription of an array of genes including apoptotic genes in addition to cell cycle, anti-oxidant and autophagy genes depending on the cellular context ^20-24^. In fact, a gradual increase in Foxo3 expression, nuclear localization, and activity during erythroid maturation balances ROS levels in early versus late erythroblasts ^15,25^. In early erythroid precursors, Foxo3 regulates erythroid cell cycling and differentiation. In late erythroblasts, Foxo3 is critical for terminal erythroid maturation, coordinated mitochondrial removal and enucleation ^15,19,25-27^. Foxo3 regulation of anti-oxidant genes in erythroid precursor cells, maintains RBCs’ half-life ^15^. Foxo3 loss in erythroid cells is balanced by the activation of TP53 with which Foxo3 shares many targets and functions ^15,28-30^. Foxo3’s steady upregulation with maturation of primary mouse erythroblasts is not associated with apoptosis ^15^. However, whether Foxo3 regulates apoptosis in the context of β-thalassemia erythropoiesis is unknown especially since depending on cellular state, Foxo3 may inhibit cell cycle and promote cell cycle exit, induce apoptosis or trigger differentiation ^31-33^. The cyclin-dependent kinase inhibitor p21 (Cdkn1a) is a direct target of both Foxo3 and p53 ^34,35^ known to repress cell cycle entry in erythroid cells ^15,36,37^ but its function in β-thalassemia erythroid cells is largely unknown ^38^.

Here, we asked whether Foxo3 is implicated in defective β-thalassemia red blood cell production and subsequent abnormalities. We find that both Foxo3 and TP53 are prematurely activated in β-thalassemia erythroid compartment. We further show using genetic approaches that loss of either Foxo3 or its target p21 reduces significantly β-thalassemia erythroid cell apoptosis. However, while Foxo3 loss improves anemia, it does not reduce splenomegaly or reticulocytes in β-thalassemic mice; on the other hand, loss of p21 does not improve β-thalassemia anemia, ineffective erythropoiesis, extramedullary erythropoiesis or splenomegaly. Altogther, these intriguing results suggest that mechanisms that control anemia, erythroid apoptosis and ineffective erythropoiesis may be uncoupled in β-thalassemia. Our findings also uncover a tight and intricate network between p21, Foxo3 and TP53 in controlling β-thalassemia erythroid apoptosis.

## Results

### Apoptotic phenotype in mouse β-thalassemia is similar to human β-thalassemia

To address the underlying mechanism of apoptosis of β-thalassemic erythroid cells, we used *Hbb*^*th3/+*^ mice that exhibits a phenotype similar to the human β-thalassemia intermedia and harbors deletions in both β-globin subunits ^39,40^. As anticipated using flow cytometry parameters of size, CD44, and TER119 markers ^41^ to analyze erythroid cell maturation, we found the heterozygous *Hbb*^*th3/+*^ erythroblasts ^39,40^ to show a delay/impairment in their maturation with a specific increase in relatively mature polychromatophilic cells found in Gate III and a corresponding decrease in Gate V red blood cells (RBCs) (**Figures 1A and 1B**). Like in β-thalassemic patients, *Hbb*^*th3/+*^ mice display splenomegaly, an indication of extramedullary erythropoiesis associated with clearing damaged RBCs (**Figure 1C**) and elevated ROS levels suggesting increased oxidative damage in late maturing β-thalassemia erythroid cells (**Supplemental Figure S1**). Interestingly, ROS levels were lower in β-thalassemic immature erythroblasts relative to WT controls (**Supplemental Figure S1**).

**Figure 1:**
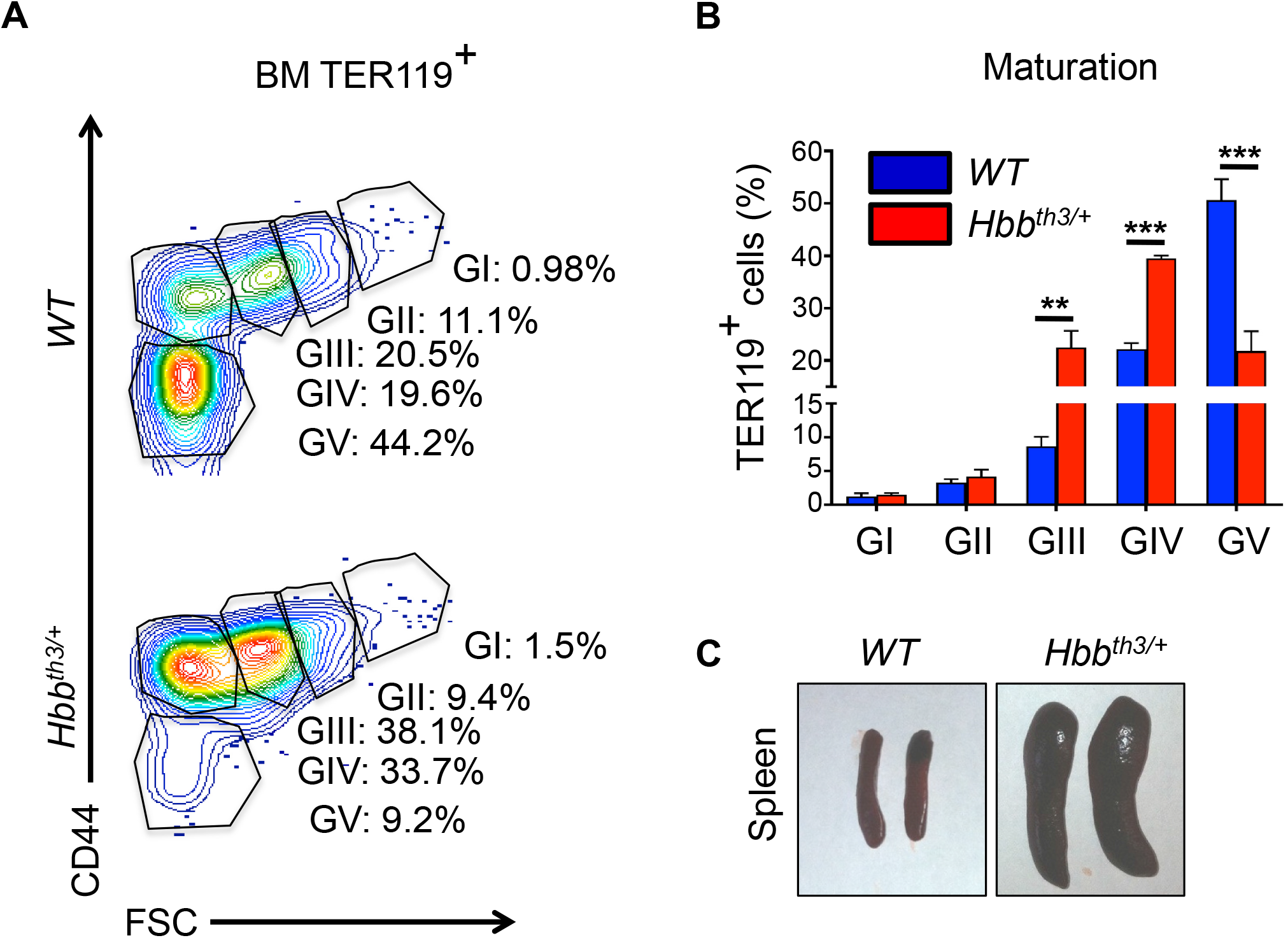
β-thalassemia mice (*Hbb*^*th3/+*^) closely mimic the intermediate form of human β-thalassemia. **A**. Gating strategy to identify erythroblasts (GI-GIII), reticulocytes (Gate IV), and RBCs (Gate V) in *WT* and *Hbb*^*th3/+*^ bone marrow cells. **B**. Histograms showing the percentage of each population in (A). Note the increased Gate III cells in *Hbb*^*th3/+*^ mice with a concomitant decrease in Gate V cells. *p < 0.05.**C**. Macroscopic appearance of spleen in *WT* and *Hbb*^*th3/+*^ mice.

To further validate the model in our own hands, we next determined that the frequency of apoptotic (Annexin V^+^ 7AAD^-^) bone marrow erythroblasts is significantly higher in *Hbb*^*th3/+*^ relative to wild type (*WT*) mice, and is best observed in relatively mature polychromatophilic erythroblasts (Gate III, **Figures 2A-2B, Supplemental Figure S1B)** as reported in patients’ β-thalassemia ^5^.

**Figure 2:**
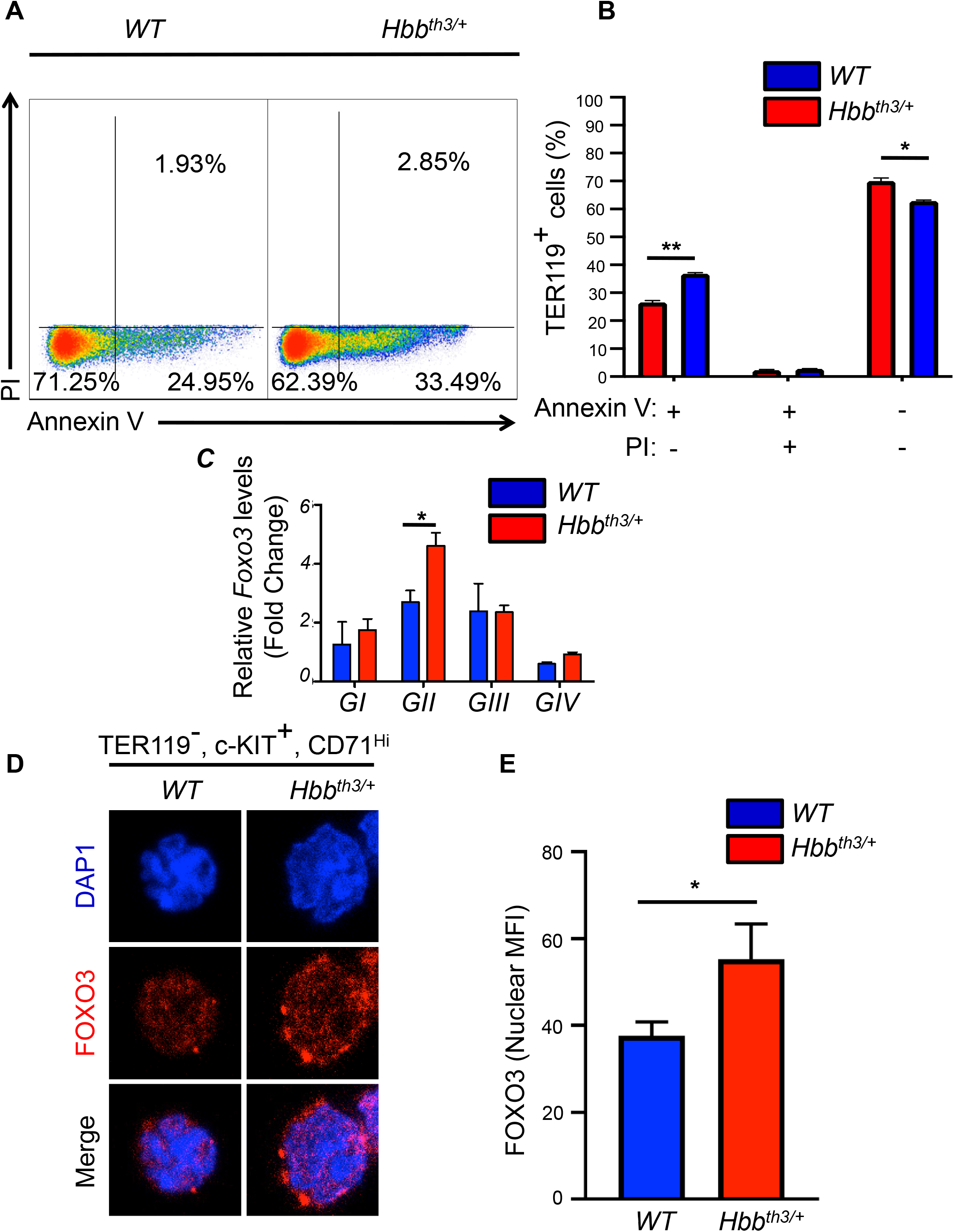
Apoptotic phenotype in mouse β-thalassemia is similar to human β-thalassemia. **A**. Annexin V/PI staining of TER119^+^ subpopulation within bone marrow isolated from *WT* and *Hbb*^*th3/+*^ mice. **B**. Quantification of apoptotic, dead, and live TER119 cells isolated from *WT* and *Hbb*^*th3/+*^ mice; *p < 0.05. **C**. qRT-PCR expression analysis of Foxo3 in gates I-IV; *p < 0.05. **D**. Foxo3 localization in erythroid progenitors using confocal microscopy. **E**. Quantification of data in (**D**); analyses of at leasy 40 cells, *p < 0.05 (n = 3).

### Foxo3 is activated prematurely mediating stress response in β-thalassemic erythroid compartment

To evaluate whether Foxo3 is implicated in regulating β-thalassemic erythroid apoptosis, its expression and nuclear localization - an indication of Foxo3 activity - was analyzed in erythroid progenitors/early erythroblasts prior to exhibiting apoptosis. Foxo3 transcript was elevated in *Hbb*^*th3/+*^ relative to *WT* early erythroblasts (**Figure 2C)**. Foxo3 protein expression and nuclear localization was also enhanced in both maturing erythroid progenitors (TER119^-^, c-KIT^+^, CD71^Hi^) ^42^ and erythroid precursor TER119^Lo^ CD71^Hi^ cells ^43^ as measured by immunofluorescence (IF) analysis (**Figures 2D-2E, Supplemental Figure S2A**). This increase was independent of any significant change in *Foxo1* mRNA expression (**Supplemental Figure S2B**) suggesting specific Foxo3 effects and together indicating that Foxo3 is activated precociously in β-thalassemic erythroid progenitors.

Since the pro-apoptotic TP53 is in a regulatory network with Foxo3 ^29,44^, we examined the TP53 status in *Hbb*^*th3/+*^ erythroblasts. qRT-PCR analysis showed increased TP53 transcript levels in erythroid precursors (**Figure 3A**). Immunofluorescence staining further confirmed the TP53 protein expression is increased in *Hbb*^*th3/+*^ TER119^-^, c-KIT^+^, CD71^Hi^ enriched for erythroid progenitors ^42^. Specifically, nuclear TP53 levels were increased in *Hbb*^*th3/+*^ TER119^-^, c-KIT^+^, CD71^Hi^ relative to the wild type controls (**Figures 3B and 3C, Supplemental Figure S2C**). These studies together suggest that overall, increased TP53/Foxo3 may be mediating the increased apoptosis in *Hbb*^*th3/+*^ erythroblasts.

**Figure 3:**
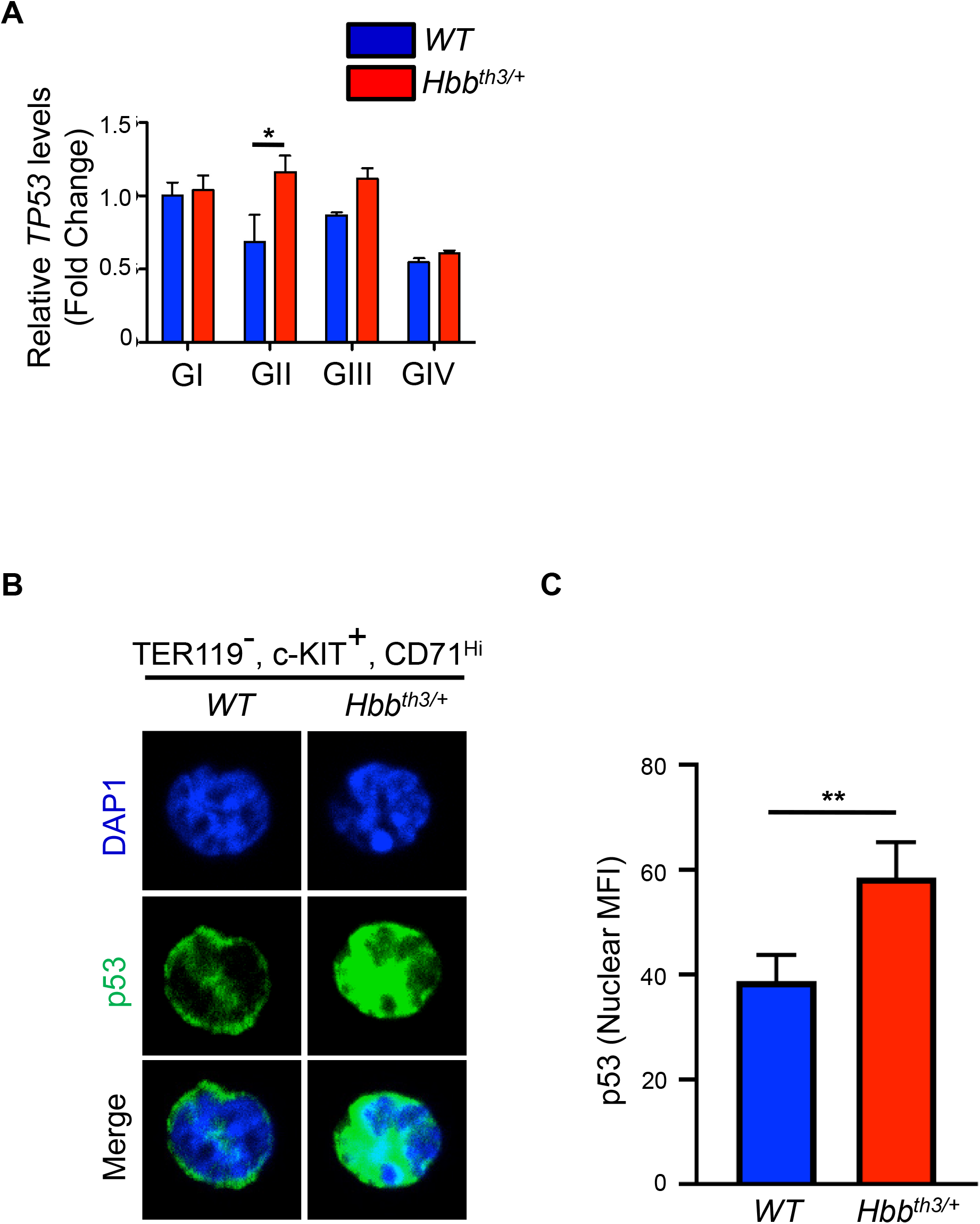
Increased p53 expression in β-thalassemic erythroid progenitors. **A**. qRT-PCR expression analysis of TP53 in gates I-IV. *p < 0.05. **B**. p53 localization in erythroid progenitors and precursors using confocal microscopy. **C**. Quantification of data in (A); analyses of at leasy 40 cells *p < 0.05 (n = 3).

### Loss of Foxo3 reduces erythroblast apoptosis and improves red blood cell production in a β-thalassemia mouse model

To further dissect the role of Foxo3 in the pathophysiology of β-thalassemia, we generated double mutant *Foxo3*^*-/-*^*/Hbb*^*th3/+*^ mice. In agreement with Foxo3 mediation of erythroid cell apoptosis, loss of Foxo3 in *Hbb*^*th3/+*^ mice improved the anemia (**Figure 4A and Table 1**) as we observed an increase in RBC production by ∼10% and hemoglobin content (by 1g/dL) (**Figure Table 1**). These significant results were notable specifically in the context of β-thalassemia. Importantly, apoptosis was markedly reduced in *Foxo3*^*-/-*^*/Hbb*^*th3/+*^ erythroblasts, specifically in Gates II - IV (**Figures 4B**,, **Supplemental Figure S3**). These results suggest that Foxo3 upregulation is pro-apoptotic in β-thalassemic erythroid compartment. However, loss of Foxo3 did not rescue the splenomegaly in *Hbb*^*th3/+*^ mice, indicating persistent extramedullary hematopoiesis (**Figure 4C**). Not surprisingly, given the Foxo3 anti-oxidant functions ^15,28^, ROS levels were increased in double mutant erythroblasts (**Supplemental Figure S4**) suggesting that improved erythroblast survival in *Foxo3*^*-/-*^*/Hbb*^*th3/+*^ mice was despite increased ROS in both bone marrow and spleen of double mutant erythroblasts (**Supplemental Figure S4**). These results indicate that the alleviation of the apoptotic phenotype in *Foxo3*^*-/-*^*/Hbb*^*th3/+*^ erythroblasts (**Figure 4; Supplemental Figure S3**) is likely acting via an oxidative stress-independent pathway.

**Table 1:** Blood parameters in *Hbb*^*th3/+*^ versus *Foxo3*^*-/-*^*/Hbb*^*th3/+*^

| Parameters | *WT* | *Foxo3<sup>-/-</sup>* | *Hbb<sup>th3/+</sup>* | *Foxo3<sup>-/-</sup>/Hbb<sup>th3/+</sup>* |
| --- | --- | --- | --- | --- |
| **RBC (M/μL)** | 10.4 ± 0.3 | 9.0 ± 0.1* | 6.7 ± 0.7* | 7.5 ± 0.6<sup>#</sup> |
| **HGB (g/dL)** | 15.0 ± 0.3 | 14.8 ± 0.1 | 8.2 ± 0.3* | 9.2 ± 0.7<sup>#</sup> |
| **HCT (%)** | 54.6 ± 1.6 | 52.8 ± 0.9 | 29.3 ± 5.2* | 34.5 ± 3.9<sup>#</sup> |
| **MCV (fL)** | 52.8 ± 1.2 | 58.5 ± 0.8* | 41.2 ± 1.4* | 45.1 ± 3.0<sup>#</sup> |
| **MCH (pg)** | 14.6 ± 0.2 | 16.4 ± 0.2 | 12.7 ± 1.6* | 12.5 ± 0.4 |
| **MCHC (g/dL)** | 27.4 ± 0.5 | 28.0 ± 0.3 | 30.2 ± 4.1 | 27.2 ± 1.4<sup>#</sup> |
| **Retic (%)** | 3.6 ± 0.3 | 6.6 ± 0.5* | 24.0 ± 5.7* | 28.3 ± 7.0 |
| **n (mice)** | 9 | 10 | 7 | 10 |

\*p < 0.05 between *WT* and *Foxo3<sup>-/-</sup>* or *Hbb<sup>th3/+</sup>*; <sup>#</sup>p < 0.05 between *Hbb<sup>th3/+</sup>* and *Foxo3<sup>-/-</sup>/Hbb<sup>th3/+</sup>*
RBC, Red Blood Cells; HCT, Hematocrit; MCV, Mean Corpuscular Volume; MCH, Mean Corpuscular Hemoglobin; MCHC, MCH Concentration, Retic, Reticulocytes

**Figure 4:**
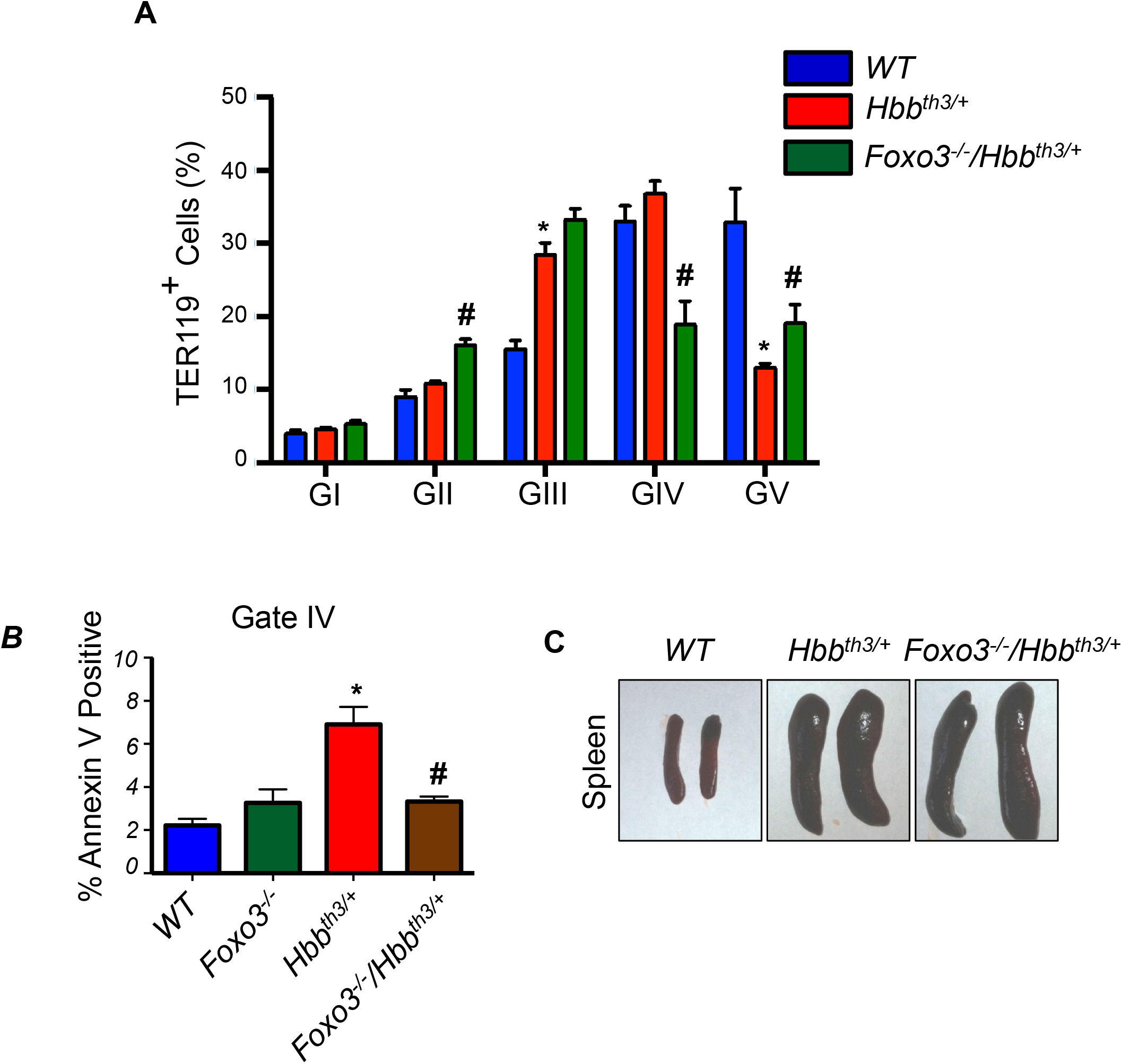
**A**. Foxo3 ablation alters BM erythroblast frequencies. *p < 0.05. **B**. Bone marrow cells from *WT, Foxo3*^*-/-*^, *Hbb*^*th3/+*^ and *Foxo3*^*-/-*^*/Hbb*^*th3/+*^ mice were isolated, FACS sorted, and stained with Annexin V to check for apoptosis. *p < 0.05 between *WT* and *Hbb*^*th3/+*^ groups and #p < 0.05 between *Hbb*^*th3/+*^ and *Foxo3*^*-/-*^*/Hbb*^*th3/+*^ groups. **C**. Macroscopic examination of spleen from *WT, Hbb*^*th3/+*^, *Foxo3*^*-/-*^*/Hbb*^*th3/+*^.

Earlier studies had shown Foxo3 to be key in regulating cell cycle in addition to apoptosis in a context- and stress-dependent manner ^15,16,18,19,25,28,45-47^. RNA-seq analyses of *Foxo3*^*-/-*^ erythroblasts ^14^ revealed deregulated expression of many apoptotic genes relative to wild type controls (**Supplemental Figure S5A**). Further, using a Fluidigm Platform we interrogated the expression of a panel of apoptotic genes by qRT-PCR analysis in double mutant *Foxo3*^*-/-*^*/Hbb*^*th3/+*^ erythroblasts. Among these we found that the expression of BCL2 family member PUMA transcript and protein (P53 up-regulated modulator of apoptosis or BCL2-binding component 3, BBC3), but not BIM, both targets of Foxo3 and TP53 ^48,49^, was greatly augmented in *Hbb*^*th3/+*^ erythroblasts (**Supplemental Figures S5B and S5C**). Notably, PUMA’s expression was significantly downregulated in double mutant *Foxo3*^*-/-*^*/Hbb*^*th3/+*^ erythroblasts (**Supplemental Figure S5B**). However, generating double mutant *Puma*^*-/-*^*/Hbb*^*th3/+*^ erythroblasts was not productive likely due to the linkage of *Puma* and *β-globin* genes on mouse chromosome 7.

### p21 ablation in mouse β-thalassemia improves apoptosis without significantly improving RBC formation

We next focused on p21 ^50^ that is another common target of both Foxo3 and TP53 ^29^. We found that *p21* transcript is highly and significantly upregulated in *Hbb*^*th3/+*^ erythroblasts throughout maturation (**Figure 5A**). The p21 protein expression was greatly enhanced in mouse *Hbb*^*th3/+*^ erythroblasts (**Figure 5B**). Importantly, expression of p21 protein was also significantly elevated in *in vitro* CD34^+^-differentiated human erythroblasts from two independent β-thalassemic patients, strongly suggesting that as in the mouse, p21 is upregulated in human β-thalassemic erythroblasts (**Figures 5C and 5D**).

**Figure 5:**
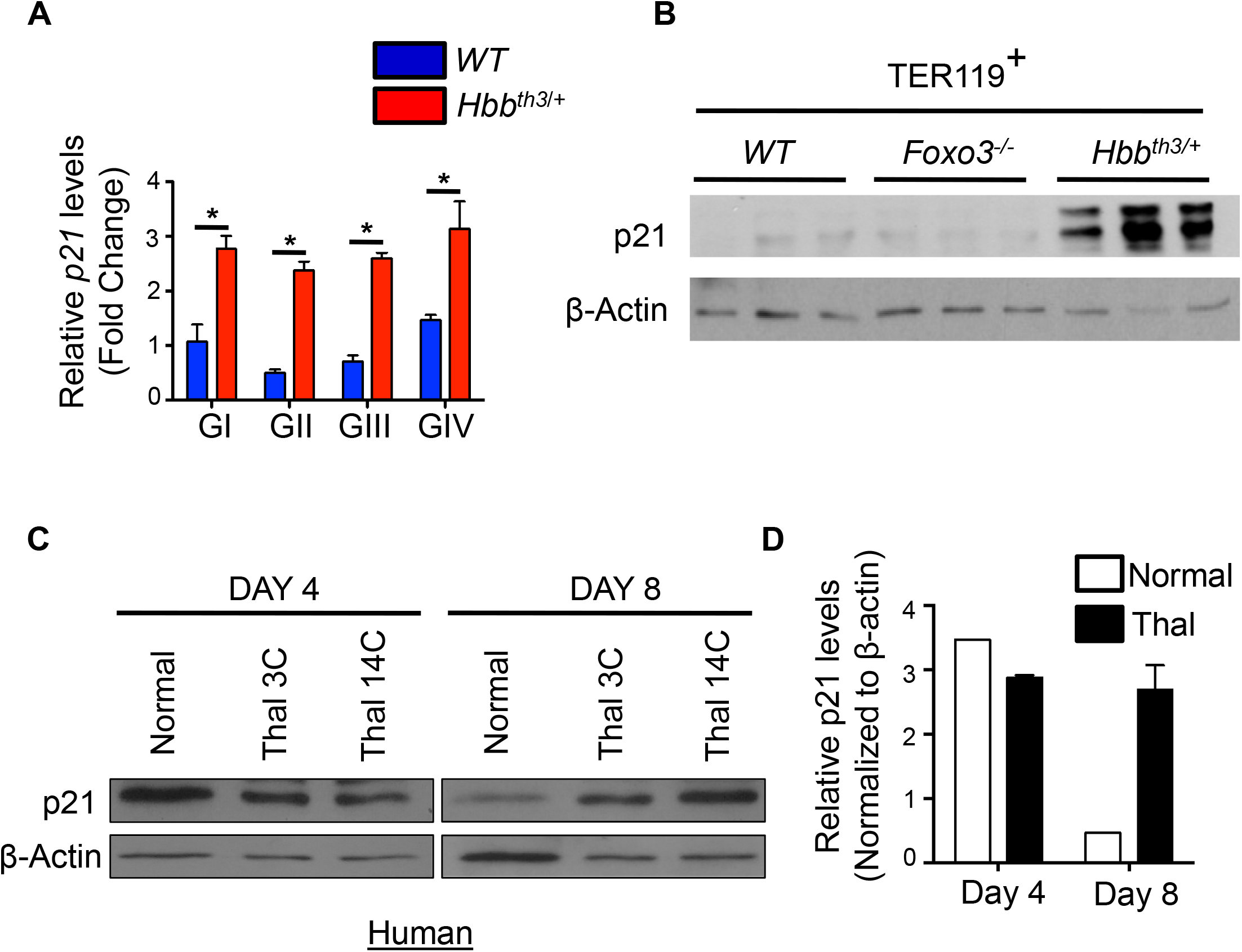
**A**. qRT-PCR expression analysis of p21. *p < 0.05. **B**. Immunoblotting showing p21 expression in *WT, Foxo3*^*-/-*^, and *Hbb*^*th3/+*^ TER119^+^ erythroblasts. **C**. Immunoblot analysis of p21 expression in erythroblasts derived from CD34^+^ cells isolated from normal donors and beta-thalassemic patients. **D**. Quantification of (**C**).

Given that p21 regulates apoptosis under certain stress conditions in addition to inhibiting cell cycle ^50,51^, we addressed the potential contribution of p21 to the enhanced apoptosis by generating *p21*^*-/-*^*/Hbb*^*th3/+*^ mice. We opted for this *in vivo* approach to avoid the impending artefacts of *in vitro* targeting on erythroblast maturation and apoptosis ^52^. The double mutant *p21*^*-/-*^*/Hbb*^*th3/+*^ mice were born following a non-Mendelian pattern and only a small subset survived to the adulthood. The subsequent experiments were carried using this *p21*^*-/-*^*/Hbb*^*th3/+*^ adult group. We found that deleting p21 on a β-thalassemic background significantly alleviated apoptosis in the *p21*^*-/-*^*/Hbb*^*th3/+*^ relative to *Hbb*^*th3/+*^ bone marrow erythroblasts (**Figure 6A; n = 6)** and to a lesser extent in the spleen erythroblasts (**Figure 6B**), suggesting that indeed upregulated p21 mediates apoptosis in the β-thalassemic erythroblasts. To our surprise, this was associated with significantly reduced cycling of *p21*^*-/-*^*/Hbb*^*th3/+*^ relative to *Hbb*^*th3/+*^ erythroblasts as evidenced by Ki67 cell cycle marker (**Figure 6C**). Combined with decreased cell cycling, ROS levels were reduced in the *p21*^*-/-*^*/Hbb*^*th3/+*^ relative to *Hbb*^*th3/+*^ bone marrow erythroblasts and almost normalized (**Figure 6D**). Unexpectedly however, and despite lack of bone marrow apoptosis and improved erythroblast survival, loss of p21 did not have any detectable effect on erythroid cell production (**Figures S6A-S6G**), RBC numbers, reticulocytes and related parameters in *p21*^*-/-*^*/Hbb*^*th3/+*^ relative to *Hbb*^*th3/+*^ mice (**Table 2**). Although loss of p21 led to reduced Epo release that signals an improved state of *Hbb*^*th3/+*^ erythropoiesis, it did not alleviate the splenomegaly, extramedullary erythropoiesis (**Figures S6A, S6B**), or ROS levels in β-thalassemic spleen erythroblasts (**Supplemental Figures S6A-S6I, S7, and Table 2**). Altogether these results suggest that reducing apoptosis in β-thalassemic erythroblasts does not improve RBC production or attenuate ineffective erythropoiesis.

**Table 2:** Blood parameters in *Hbb*^*th3/+*^ versus *p21*^*-/-*^*/Hbb*^*th3/+*^

|  | *WT* | *p21<sup>-/-</sup>* | *Hbb<sup>th3/+</sup>* | *p21<sup>-/-</sup>/Hbb<sup>th3/+</sup>* |
| --- | --- | --- | --- | --- |
| **RBC (M/μL)** | 10.9 ± 0.7 | 10.5 ± 0.3 | 8.4 ± 0.5 | 8.2 ± 0.4 |
| **HGB (g/dL)** | 15.6 ± 0.9 | 15.3 ± 0.3 | 8.9 ± 0.5 | 8.8 ± 0.4 |
| **WBC (K/μL)** | 3.3 ± 1.0 | 5.3 ± 1.9 | 13.3 ± 7.0 | 9.0 ± 3.9* |
| **Lympho (K/μL)** | 2.6 ± 0.7 | 3.6 ± 1.9 | 9.3 ± 5.3 | 6.7 ± 2.9 |
| **EO (K/μL)** | 0.04 ± 0.02 | 0.11 ± 0.09 | 0.24 ± 0.19 | 0.18 ± 0.17 |
| **Retic (%)** | 3.6 ± 0.4 | 3.6 ± 0.5 | 27.9 ± 2.0 | 28 ± 4.1 |
| **n (mice)** | 6 | 6 | 6 | 6 |

\*p < 0.05 between *Hbb<sup>th3/+</sup>* and *p21<sup>-/-</sup>/Hbb<sup>th3/+</sup>*
RBC, Red Blood Cells; HGB, Hemoglobin; WBC, White Blood Cells; Lympho, Lymphocytes; EO, Eosinophils, Retic, Reticulocytes

**Figure 6:**
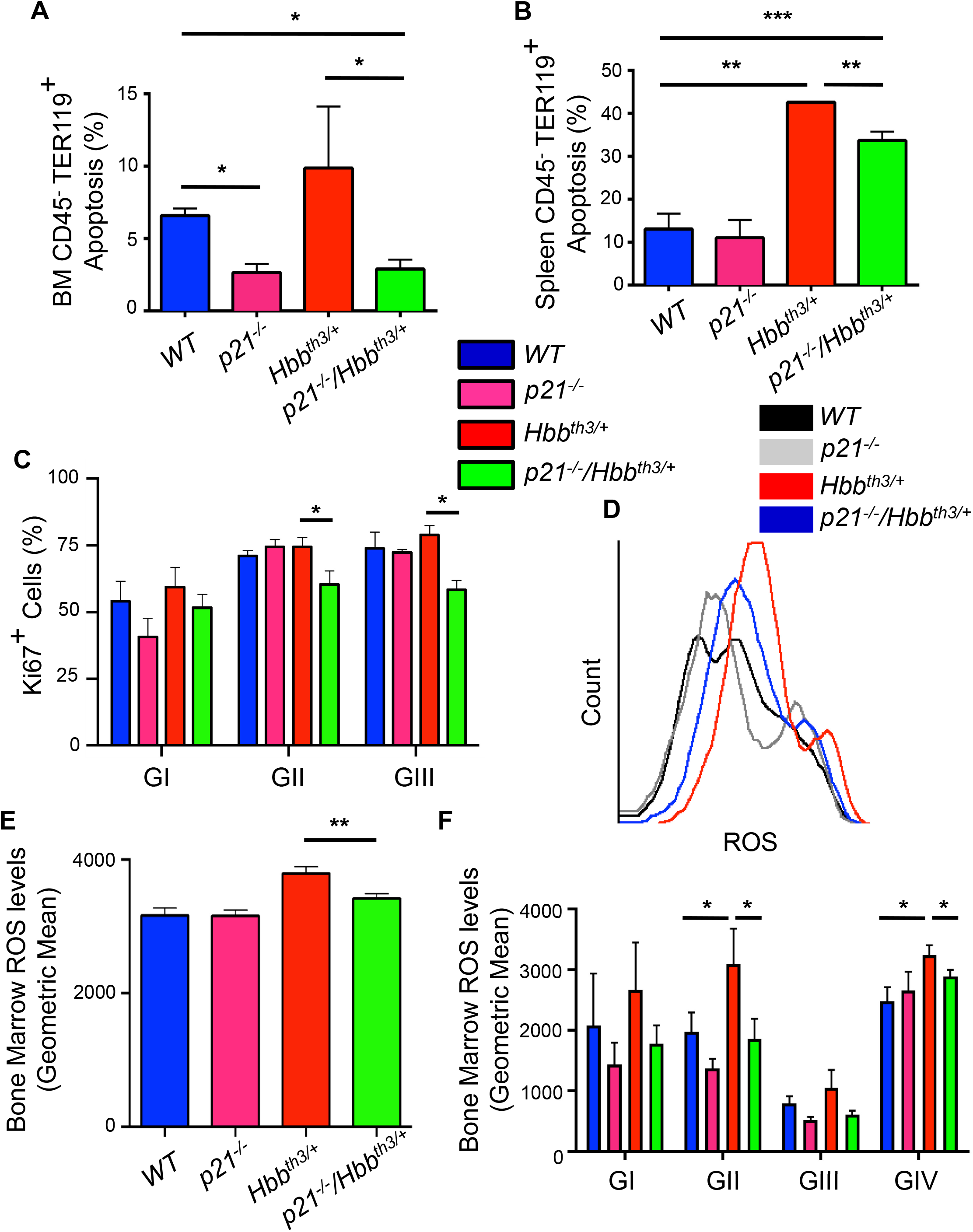
**A**. Flow cytometric analysis of apoptosis in bone marrow erythroid cells. The cells were stained with Annexin V and 7AAD to sort the TER119^+^ cells in bone marrow from each group of mice (n = 3 for each group). Each bar represents mean ± SD. *p < 0.05 between *Hbb*^*th3/+*^ and *p21*^*-/-*^*/Hbb*^*th3/+*^ and between *p21*^*-/-*^*/Hbb*^*th3/+*^ and *WT* groups. **B**. Flow cytometric analysis of apoptosis in splenic erythroid cells. The cells were stained with Annexin V and 7AAD to sort the TER119^+^ cells in spleens isolated from each group of mice (n = 3 for each group). Each bar represents mean ± SD. *p < 0.05 between *Hbb*^*th3/+*^ and *p21*^*-/-*^*/Hbb*^*th3/+*^ and between *p21*^*-/-*^*/Hbb*^*th3/+*^ and *WT* groups. **C**. Flow cytometric analysis of cell proliferation in bone marrow erythroid cells. TER119^+^ cells in the bone marrow from each group of mice (n = 3 for each group) were stained with Ki67. Each bar represents mean ± SD. *p < 0.05 between *Hbb*^*th3/+*^ and *p21*^*-/-*^*/Hbb*^*th3/+*^groups. **D**. Histogram of ROS levels in bone marrow erythroblasts as measured by DCF probe **E**. Quantification of D (n = 3), Mean ± SD **p < 0.01 between *Hbb*^*th3/+*^ and *p21*^*-/-*^*/Hbb*^*th3/+*^ groups. **F**. ROS levels in erythroblast populations obtained from each group of mice within gates I-V. Mean ± SD *p < 0.05 between p21^-/-^, *Hbb*^*th3/+*^ and *p21*^*-/-*^*/Hbb*^*th3/+*^ groups.

Decreased cycling of *p21*^*-/-*^*/Hbb*^*th3/+*^ relative to *Hbb*^*th3/+*^ erythroblasts was surprising. To have an insight into mechanisms that regulate *p21*^*-/-*^*/Hbb*^*th3/+*^ cycling, we analyzed *p21*^*-/-*^*/Hbb*^*th3/+*^ erythroblasts by confocal microscopy IF staining for the expression of Foxo3 and TP53 as these proteins together regulate erythroid cell cycling ^15^ (**Figures 7A and 7B**). While Foxo3 nuclear expression was substantially enhanced in *p21*^*-/-*^*/Hbb*^*th3/+*^ TER119^-^, c-KIT^+^, CD71^Hi^ cells, an indication of Foxo3 activation in erythroblast precursors (**Figure 7A**), we noticed that the nuclear expression of TP53 protein had plummeted specifically in these double mutant cells while significantly augmented in *Hbb*^*th3/+*^ and *p21*^*-/-*^ relative to *WT* controls (**Figure 7B**). There was no difference in the overall protein expression of Foxo3 (Figure S8, Top right), while TP53 expression was decreased in *Hbb*^*th3/+*^, *p21*^*-/-*^ and *p21*^*-/-*^*/Hbb*^*th3/+*^ erythroid cells (**Figure S8, bottom right**). These studies suggest that upregulated Foxo3 is likely mediating the p21-independent cell cycle inhibition in *p21*^*-/-*^*/Hbb*^*th3/+*^ erythroblasts. In addition, these results identify the interplay between p21, Foxo3, and TP53 in regulating *Hbb*^*th3/+*^ erythroblasts’ cycling and apoptosis. To have an insight into the reduced ROS levels in *p21*^*-/-*^*/Hbb*^*th3/+*^ erythroblasts, and considering that Foxo3 is implicated in regulating mitochondria ^53-55^ and ROS levels ^15,28,46,56,57^ we analyzed mitochondrial shape that modifies, and responds to, ROS levels and cell cycle ^58-60^. We found that mitochondria are significantly fragmented in *Hbb*^*th3/+*^ *vs*. WT erythroblasts; in addition, mitochondrial fragmentation was markedly increased in *p21*^*-/-*^*/Hbb*^*th3/+*^ even as compared to *Hbb*^*th3/+*^ erythroblasts, especially in Gate 3 (**Figure 7C**). These results were intriguing since mitochondrial fragmentation is often associated with apoptosis; and might explain reduced cell cycling. Thus, increased mitochondrial fragmentation is associated with cell cycle inhibition, reduced ROS levels, and decreased apoptosis in *p21*^*-/-*^*/Hbb*^*th3/+*^ relative to *Hbb*^*th3/+*^ erythroblasts.

**Figure 7:**
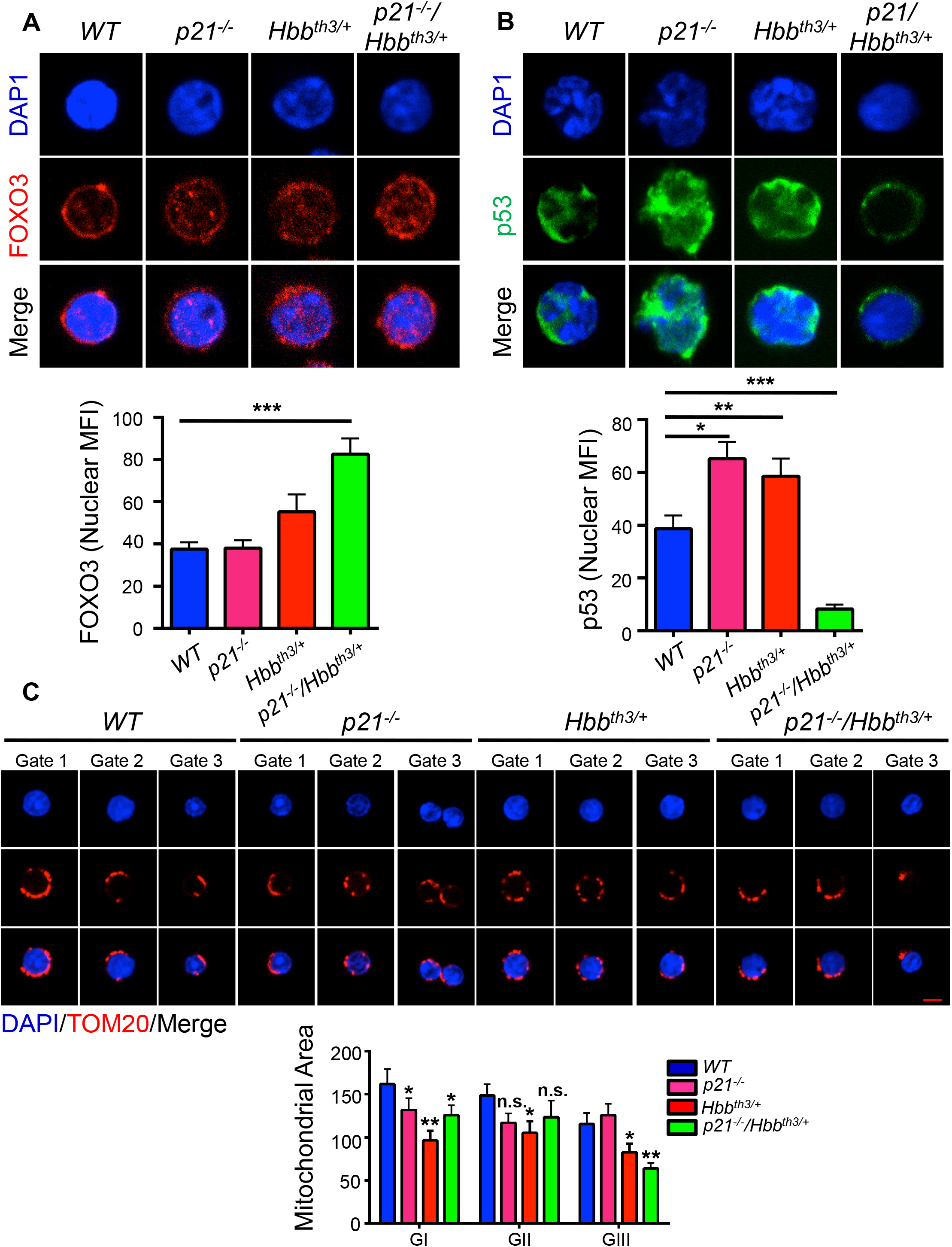
**A**. Foxo3 nuclear localization in erythroid progenitors and precursors from *WT, p21*^*-/-*^, *Hbb*^*th3/+*^ and *p21*^*-/-*^*/Hbb*^*th3/+*^ mice using confocal microscopy. ***p < 0.001. **B**. p53 nuclear localization in erythroid progenitors and precursors from *WT, p21*^*-/-*^, *Hbb*^*th3/+*^ and *p21*^*-/-*^*/Hbb*^*th3/+*^ mice using confocal microscopy. Analyses of at least 40 cells *p < 0.05, **p < 0.01, ***p < 0.001. **C**. Mitochondrial morphology in erythroid progenitors and precursors from *WT, p21*^*-/-*^, *Hbb*^*th3/+*^ and *p21*^*-/-*^*/Hbb*^*th3/+*^ mice using confocal microscopy; analyses of at least 30 cells. Mean ± S.E.M *p < 0.05, **p < 0.01. n.s. (not significant). Scale Bar: 5 µm

## Discussion

Here we found that p21 (Cdkn1a) expression is upregulated in both mouse and human β-thalassemic erythroid cells; we further showed that the elevated p21 mediates apoptosis in adult β-thalassemic *Hbb*^*th3/+*^ bone marrow and to a much lesser extent spleen erythroid cells. Notably, given that loss of p21 is not lethal in mice ^61,62^, the in-utero death of *p21*^*-/-*^*/Hbb*^*th3/+*^ mice suggests that p21 expression is necessary for the survival of β-thalassemic *Hbb*^*th3/+*^ mice. Our results indicate that paradoxically the greatly enhanced p21 expression, supports β-thalassemic erythroid precursor cell cycling (**Fig. 6C**). p21 functions in a molecular network with Foxo3 and TP53, that are both transcriptional regulators of cell cycle inhibition and apoptosis ^45,63-66^. We also showed that reducing apoptosis and significantly improving *Hbb*^*th3/+*^ erythroblasts’ survival do not ameliorate RBC production (in *p21*^*-/-*^*/Hbb*^*th3/+*^ mice) nor reduce the extramedullary erythropoiesis in β-thalassemic *Hbb*^*th3/+*^ mouse model [*p21*^*-/-*^*/Hbb*^*th3/+*^ or *Foxo3*^*-/-*^*/Hbb*^*th3/+*^ mice]. The combination of reduced apoptosis and erythroid cell division results in no gain in RBC numbers or reduction of reticulocytes in *p21*^*-/-*^*/Hbb*^*th3/+*^ mice. Notably loss of p21 in *Hbb*^*th3/+*^ mouse had much lesser effect on the splenic erythroid cells, including spleen apoptosis (reduction by >75% in apoptosis of BM CD45-TER119+ cells *vs*. < 25% in the spleen in *p21*^*-/-*^*/Hbb*^*th3/+*^ vs. *Hbb*^*th3/+*^ mice), erythroid cell numbers, or ROS levels. These were despite reducing Epo levels, suggesting distinct regulation of β-thalassemic erythroid cell maturation in the spleen *vs*. BM. Reduced ROS levels in the context of loss of p21 was not associated with improved β-thalassemia RBC production or erythroblast survival in the spleen. These findings indicate that although increased apoptosis is associated with the expansion of erythroblast pool in this β-thalassemic *Hbb*^*th3/+*^ mouse model, it is likely that apoptosis is not the driver of upstream β-thalassemic erythroid response in cell expansion. Altogether these findings raise the possibility that ineffective erythropoiesis and apoptosis in β-thalassemia may be uncoupled, at least in mice. Specifically, reduced/elimination of apoptosis in the bone marrow does not affect significantly extramedullary hematopoiesis or splenomegaly. These findings also suggest that the spleen *vs*. bone marrow microenvironment is critical in regulating β-thalassemic erythroid cell survival.

The cyclin-dependent kinase inhibitor p21 is mostly known for its function in cell cycle arrest through regulation of cyclin-dependent kinase (CDK) proteins ^50^. In addition, many complex functions are attributed to p21 including both pro- and anti-apoptotic properties ^50,67-69^. The pro-apoptotic p21 function is p53-dependent or -independent; mediators of p21-proapoptotic are poorly defined ^66^. Our genetic studies identify pro-apoptotic functions for p21 in the context of β-thalassemic erythropoiesis, they also provide evidence supporting the notion that p21 mediates apoptosis in a p53-independent manner.

Loss of Foxo3 clearly improved red blood cell production in *Hbb*^*th3/+*^ β-thalassemic mice. This improvement was significant in the context of β-thalassemia as a gain of 1g/dL of hemoglobin (as in *Foxo3*^*-/-*^*/Hbb*^*th3/+*^ erythroblasts), equivalent to one unit of blood, may notably reduce the transfusion schedule in β-thalassemic patients. Given the erythroid phenotype in *Foxo3*^*-/-*^ mice ^15,25^, these results were unanticipated. We had previously found that loss of Foxo3 results in increased circulating Epo, ROS-mediated constitutive activation of both protein kinases JAK2 and mTORC1 that sustain Epo-receptor (EpoR) signaling and cell growth ^25^. In addition, we had shown that inhibiting JAK2 ^38^ or mTORC1 ^25^ improves RBC production and reduces anemia in *Hbb*^*th3/+*^ β-thalassemic mice. However, decreased apoptosis, increased survival of *Foxo3*^*-/-*^*/Hbb*^*th3/+*^ erythroblasts (**Figures 4B, Supplemental Figure S3**) and improved RBC production (**Table 1, Figure 4B**) had no impact on reticulocytosis or splenomegaly suggesting lack of amelioration of ineffective erythropoiesis in double mutant mice. Given the importance of Foxo3 in RBC production and reticulocyte maturation ^15,17^, including mitochondrial removal and enucleation ^14,25^, it is also unclear whether the quality of RBC formed in *Foxo3*^*-/-*^*/Hbb*^*th3/+*^ mice is optimal or whether potential therapies targeting Foxo3 be beneficial for β-thalassemia RBC production.

Our findings indicate that a network of p21, Foxo3 and TP53 coordinates erythroid cell cycling and apoptosis in β-thalassemia mice in an unanticipated fashion presumably to balance cell cycle and apoptosis. Our studies also expose the distinct regulation of β-thalassemic erythroid cell maturation in the spleen *vs*. bone marrow.

## Supporting information

Revised Supplemental Files

## Acknowledgments

The authors thank the Flow Cytometry Core and Microscopy CoRE at the Icahn School of Medicine at Mount Sinai for technical help, Dr. Miriam Merad for providing p21^-/-^ mice and Dr. Xiuli An (New York Blood Center) for providing anti-human erythroid antibodies. Work in SR’s lab is supported by Commonwealth Universal Research Enhancement (C.U.R.E.) Program Pennsylvania, CuRED-Frontier Program, and NIH NIDDK (R01 DK090554, R01 DK095112). This work was supported by R01HL136255, R01CA205975 and funds from NYSTEM IIRP C32602GG to SG.

COI: S.R. is a member of scientific advisory board of Ionis Pharmaceuticals, Meira GTx, Incyte and Disc Medicine and owns stock options from Disc Medicine. S.R. has been or is consultant for Cambridge Healthcare Res, Celgene Corporation, Catenion, First Manhattan Co., FORMA Therapeutics, Ghost Tree Capital, Keros Therapeutics, Noble insight, Protagonist Therapeutics, Sanofi Aventis U.S., Slingshot Insight, Techspert.io and BVF Partners L.P., Rallybio, LLC, venBio Select LLC.

## Materials and Methods

### Mice

All protocols were approved by the Institutional Animal Care and Use Committee of Mount Sinai School of Medicine. *Foxo3*^*+/-*^ mice were previously described ^28^. Wild-type mice were used as controls in all experiments. For all experiments, 8-12-week-old mice were used. The thalassemic mice used were previously described ^39^. The *Foxo3*^*-/-*^*/Hbb*^*th3/+*^ double mutant mice were generated by crossing the *Foxo3*^*-/-*^ with the *Hbb*^*th3/+*^ mice. *P21*^*-/-*^ mice (Jackson Laboratory) were obtained from Dr. Miriam Merad at Mount Sinai. Birth frequency of *p21*^*-/-*^*/Hbb*^*th3/+*^ deviated significantly from Mendelian inheritance (χ ^2^ chi-squared test P < 0.0001) suggesting occurance of pre-natal lethality.

### Immunoblotting

TER119^+^ cells isolated from WT, *Foxo3*^*-/-*^ and the *Hbb*^*th3/+*^ mice were boiled directly in Laemmli sample buffer at 95°C for 10min and stored at -80°C till further use. Electrophoresis was carried out and the gels were transferred onto PVDF Immobilon-P membrane (Millipore). Membranes were then blocked with 5% BSA/1XPBS/0.1% Tween-20 solution. After 3 washes, the membranes were incubated with the respective primary antibodies overnight at 4°C in 1% BSA/1XPBS/0.1% Tween-20 solution. After further washes, the membranes were then incubated with the secondary antibodies for 1hr at room temperature. The blots were then washed 3 times and developed using the ECL reagent (Pierce). CD34^+^ cells were isolated from blood obtained from normal and thalassemic patients, lysates prepared as mentioned above and probed with p21 antibody. The blots were then washed and developed using the ECL reagent.

### Plasma Erythropoietin (Epo) measurement

Plasma was collected using heparin as an anti-coagulant from blood obtained from *WT, p21*^*-/-*^, *Hbb*^*th3/+*^ and *p21*^*-/-*^*/Hbb*^*th3/+*^ mice. It was then centrifuged for 20 min at 2000g within 30 min of collection. Epo was quantified using the mouse Erythropoietin/Epo Quantikine ELISA Kit (R&D Systems), according to manufacturer’s instructions.

### Hematological Studies

Blood samples were obtained from *WT, Foxo3*^*-/-*^, *Hbb*^*th3/+*^, *p21*^*-/-*^, *Foxo3*^*-/-*^*/Hbb*^*th3/*^, and *p21*^*-/-*^*/Hbb*^*th3/+*^ mice, right after sacrificing the mice and collected in EDTA or Heparin. Complete blood counts (CBC) were measured at the Comparative Pathology lab at Mount Sinai using Advia 120 analyzer.

### Assay To Measure Intracellular ROS

TER119^+^ cells were isolated from the bone marrow of indicated mice. These cells were washed with PBS and then resuspended in PBS supplemented with 2% FCS or FBS and 5µM CM-H2DCFDA probe (Invitrogen) and incubated in the dark for 20min at 37°C under 5% CO2. The oxidative conversion of CM-H2DCFDA to the fluorescent product was measured using flow cytometry as previously described ^46,54^.

### Measurement of apoptosis using Annexin V-PI staining

Erythroblasts from the bone marrow or spleen were isolated from *WT, Foxo3*^*-/-*^, *Hbb*^*th3/+*^, *21*^*-/-*^, *Foxo3*^*-/-*^*/Hbb*^*th3/+*^, and *p21*^*-/-*^*/Hbb*^*th3/+*^ mice as previously described ^46,54^. FACS sorted erythroblasts of indicated stages were stained with TER119 and CD44 and apoptosis was analyzed by flow cytometry using the Annexin V-FITC Apoptosis Detection Kit (eBioscience).

### Ki-67 Staining

FACS purified erythroblasts of indicated stages were isolated from *WT, p21*^*-/-*^, *Hbb*^*th3/+*^ and *p21*^*-/-*^*/Hbb*^*th3/+*^ mice and stained for Ki-67 using the Mouse Anti-Ki-67 Set (BD Biosciences, Cat# 556026). Briefly, cells were fixed in 70-80% ethanol at -20°C for 2hr. These were then washed 2X at 1000rpm for 10min with PBS supplemented with 1%FBS and 0.09% sodium azide and resuspended at a concentration of a million cells/100μL. After transferring into a fresh tube, 20μL antibody was added, gently mixed, and incubated at RT for 20-30min in the dark. After 2X washes with PBS at 1000rpm for 5min, supernatant was discarded and the cells were resuspended in 500μL PBS and 10μL of PI solution was added. FACS analysis was carried out using the BD LSRII.

### Immunofluorescence Staining and Confocal Microscopy

FACS-sorted erythroid progenitors (TER119^-^, c-KIT^+^, CD71^Hi^) or TER119^Lo^ CD71^Hi^ cells from bone marrow of *WT, Hbb*^*th3/+*^, *p21*^*-/-*^, and *p21*^*-/-*^*/Hbb*^*th3/+*^ were cytospun at 250rpm for 3min. These were then fixed with 4% paraformaldehyde and permeabilized with 0.1% Triton X-100/1X PBS solution. Following blocking in 1% BSA/1X PBS solution for 1hr, the slides were incubated with the primary antibodies overnight (anti-p53 and anti-FOXO3). The slides were then washed 3 times with 1X PBS and incubated at RT for 1hr with the secondary antibody. The slides were again washed 3 times with 1X PBS, air dried, and mounted with Fluoroshield mounting medium containing DAPI (Abcam, Cat# ab104139).

### Human CD34 Purification and Culture

Peripheral blood were obtained from normal and Beta-thalassemic patients (IRB: 15-012123, CHOP, Dr. Stefano Rivella) and CD34^+^ cells were isolated by positive selection using the human CD34 Microbead kit from Miltenyi Biotec (Catalog No.130-046-702) as per manufacturer’s instructions. The CD34 culture and differentiation was performed as previously described ^27,70^. CD235a (Glycophorin A), CD49d (α4 integrin), and CD233 (Band 3) antibodies were obtained from Dr.Xiuli An’s laboratory (New York Blood Center) and used for determining the stage of erythroid cell differentiation as in ^27^.

### qRT-PCR analysis

Total RNA was isolated using the RNeasy mini kit (Qiagen). qRT-PCR analysis was carried out as described previously ^14,25^.

