## Supplementary material for "Elevated p21 (CDKN1a) mediates β-thalassemia erythroid apoptosis but its loss does not improve β-thalassemic erythropoiesis": Revised Supplemental Files

**Affiliations:** [1]Department of Cell, Developmental & Regenerative Biology, [2]Developmental and Stem Cell Biology Multidisciplinary Training, Graduate School of Biomedical Sciences, Icahn School of Medicine at Mount Sinai, New York, New York 10029, [3] Children's Hospital of Philadelphia, University of Pennsylvania, Philadelphia, PA 19104, [4]Department of Oncological Sciences, [5]Black Family Stem Cell Institute, [6]Tisch Cancer Institute, Icahn School of Medicine at Mount Sinai, New York, New York 10029

**# Correspondence:** Saghi Ghaffari, M.D.-Ph.D., Department of Cell, Developmental & Regenerative Biology, Icahn School of Medicine at Mount Sinai, New York, NY 10029, Tel: 212-659-8271, Fax: 212-803-6740;

### Key Points

- Elevated p21 mediates β-thalassemia erythroid cell apoptosis
- Loss of Foxo3 or p21 reduces β-thalassemia erythroid cell apoptosis but does not improve β-thalassemia ineffective erythropoiesis
- A network of Foxo3, p21 and TP53 controls β-thalassemia erythroid apoptosis
- Apoptosis may be uncoupled from ineffective erythropoiesis in β-thalassemia

**Running Title: p21 regulates β-thalassemia erythroid apoptosis**

§ Current position: Department of Therapeutic Radiology, Yale University, Connecticut
@ Current position: HemoGenyx Pharmaceuticals
*These authors contributed equally to the work

**Supplemental Figure Legends**

**Figure S1: A.** ROS is observed to be lower in GI, GII and GIII thalassemic erythroblasts as measured with the ROS probe DCF. **B.** Annexin V staining of *WT* and $Hbb^{th3/+}$ BM erythroblasts within gates I-V. Data as Mean ± S.E.M; *p < 0.05, **p < 0.01.

**Figure S2: A.** Foxo3 localization in erythroid progenitors and precursors using confocal microscopy. Right Panel: Quantification of data in **A.** Representative data as Mean ± S.E.M of at least 30 cells (n = 3) * p < 0.05. **B.** qRT-PCR expression analysis of *Foxo1* in gates I-IV. **C.** p53 localization in erythroid progenitors and precursors using confocal microscopy. Right Panel: Quantification of data in **C.** Representative data as Mean ± S.E.M of at least 30 cells (n = 3) * p < 0.05.

**Figure S3:** Annexin V staining of *WT*, $Foxo3^{-/-}$, $Hbb^{th3/+}$, $Foxo3^{-/-}/Hbb^{th3/+}$ BM erythroblasts within gates I-V. Representartive of at lest 3 experiments Mean ± S.E.M *p < 0.05, **p < 0.01.

**Figure S4**: **A.** ROS levels in erythroblast populations obtained from bone marrow from each group of mice. **B.** ROS levels in erythroblast populations obtained from spleen from each group of mice. Representartive of at lest 3 experiments Mean ± S.E.M *p < 0.05, **p < 0.01.

**Figure S5: A.** Heatmap of RNA-Seq data of apoptosis-related gene cluster in Gates I-III of *WT* and $Foxo3^{-/-}$ erythroblasts. **B.** qRT-PCR expression analysis of pro-apoptotic genes *Puma* and *Bim* in $Hbb^{th3/+}$ and double mutant erythroblasts. **C.** PUMA is upregulated in β-thalassemic erythroblasts. Mean ± S.E.M *p < 0.05 (n=3).

**Figure S6**: **A.** Macroscopic examination of spleen from *WT*, $p21^{-/-}$, $Hbb^{th3/+}$, $p21^{-/-}/Hbb^{th3/+}$. **B.** Mean spleen weight ± SD of *WT*, $p21^{-/-}$, $Hbb^{th3/+}$, and $p21^{-/-}/Hbb^{th3/+}$ mice from 3 mice per group. **C.** Flow cytometric analysis of splenocytes stained with the TER119 antibody. n = 3 for each group. **D.** Splenocyte count from *WT*, $p21^{-/-}$, $Hbb^{th3/+}$ and $p21^{-/-}/Hbb^{th3/+}$ mice from 3 mice per group. Bars represent mean ± SD. **E.** Total number of bone marrow cells analyzed by flow cytometry. n = 3 for each group. **F.** Total live cells within the bone marrow cells quantified in (**E**). n = 3 for each group. **G.** Flow cytometric analysis of bone marrow cells stained with the TER119 antibody. n = 3 for each group. **H.** ROS levels in erythroblast populations obtained from spleen from each group of mice (n = 3). **I.** Epo levels in erythroblast populations obtained from bone marrow from each group of mice (n = 3).

**Figure S7: A.** Histograms showing the percentage of erythroblasts (GI-GIII), reticulocytes (Gate IV), and RBCs (Gate V) in *WT*, $p21^{-/-}$, $Hbb^{th3/+}$, $p21^{-/-}/Hbb^{th3/+}$ bone marrow cells. **B.** Histograms showing the percentage of erythroblasts (GI-GIII), reticulocytes (Gate IV), and RBCs (Gate V) in *WT*, $p21^{-/-}$, $Hbb^{th3/+}$, $p21^{-/-}/Hbb^{th3/+}$ splenocytes.

**Figure S8: A.** Foxo3 and p53 nuclear localization in $TER119^{+}$ cells from *WT*, $p21^{-/-}$, $Hbb^{th3/+}$, and $p21^{-/-}/Hbb^{th3/+}$ mice using confocal microscopy. Analyses of at least 40 cells $^{*}p < 0.05$.

**Supplemental Table 1**

Primary antibodies used in flow cytometry and immunofluorescence

| ***Antibody*** | ***Source*** | ***Species*** | ***Clone*** | ***Dilution*** | ***Catalog Number*** |
|---|---|---|---|---|---|
| **FLOW CYTOMETRY** | | | | | |
| Mouse TER119 (FITC) | BD Biosciences | Rat | TER-119 | 1:100 | 557915 |
| Mouse TER119 (PE) | BD Biosciences | Rat | TER-119 | 1:100 | 553673 |
| Mouse CD44 (V450) | BD Biosciences | Rat | IM7 | 1:100 | 560451 |
| Mouse CD45 (APC) | BD Biosciences | Rat | 30-F11 | 1:100 | 559864 |
| **IMMUNOFLUORESCENCE** | | | | | |
| Tom20 | Santa Cruz | Rabbit | FL-145 | 1:100 | sc-11415 |
| p53 | Santa Cruz | Mouse | DO-1 | 1:100 | sc-126 |
| FoxO3a | Cell Signaling | Rabbit | D19A7 | 1:100 | 12829S |
| **IMMUNOBLOTTING** | | | | | |
| PUMA | proteintech | Rabbit | - | 1:200-1:500 | 55120-1-AP |
| p21 | Cell Signaling | Rabbit | 12D1 | 1:200-1:500 | 2947 |
| β-actin | Cell Signaling | Rabbit | 13E5 | 1:2000-1:5000 | 4970 |

**Supplemental Table 2**

Secondary antibodies

| ***Antibody*** | ***Source*** | ***Species*** | ***Fluorophore*** | ***Dilution*** | ***Catalog Number*** |
|---|---|---|---|---|---|
| **IMMUNOFLUORESCENCE** | | | | | |
| Anti-Rabbit | Invitrogen | Goat | Alexa Fluor 594 | 1:1000 | A11012 |
| Anti-Mouse | Invitrogen | Goat | Alexa Fluor 488 | 1:400 | A28175 |
| **IMMUNOBLOTTING** | | | | | |
| Anti-Rabbit | Jackson ImmunoResearch | Goat | - | 1:2000-1:4000 | 111-035-003 |

**Supplemental Table 3**

| ***Target Gene (murine)*** | ***Primer ID*** | ***Primer Sequence*** |
|---|---|---|
| *Puma* | Forward | 5′-CTGTATCCTGCAGCCTTTGC-3′ |
| | Reverse | 5′-ACGGGCGACTCTAAGTGCT-3′ |
| *Bim* | Forward | 5′-CGACAGTCTCAGGAGGAACC-3′ |
| | Reverse | 5′-CCTTCTCCATACCAGACGGA-3′ |

Primer sequences for qPCR analyses

Fig. S1

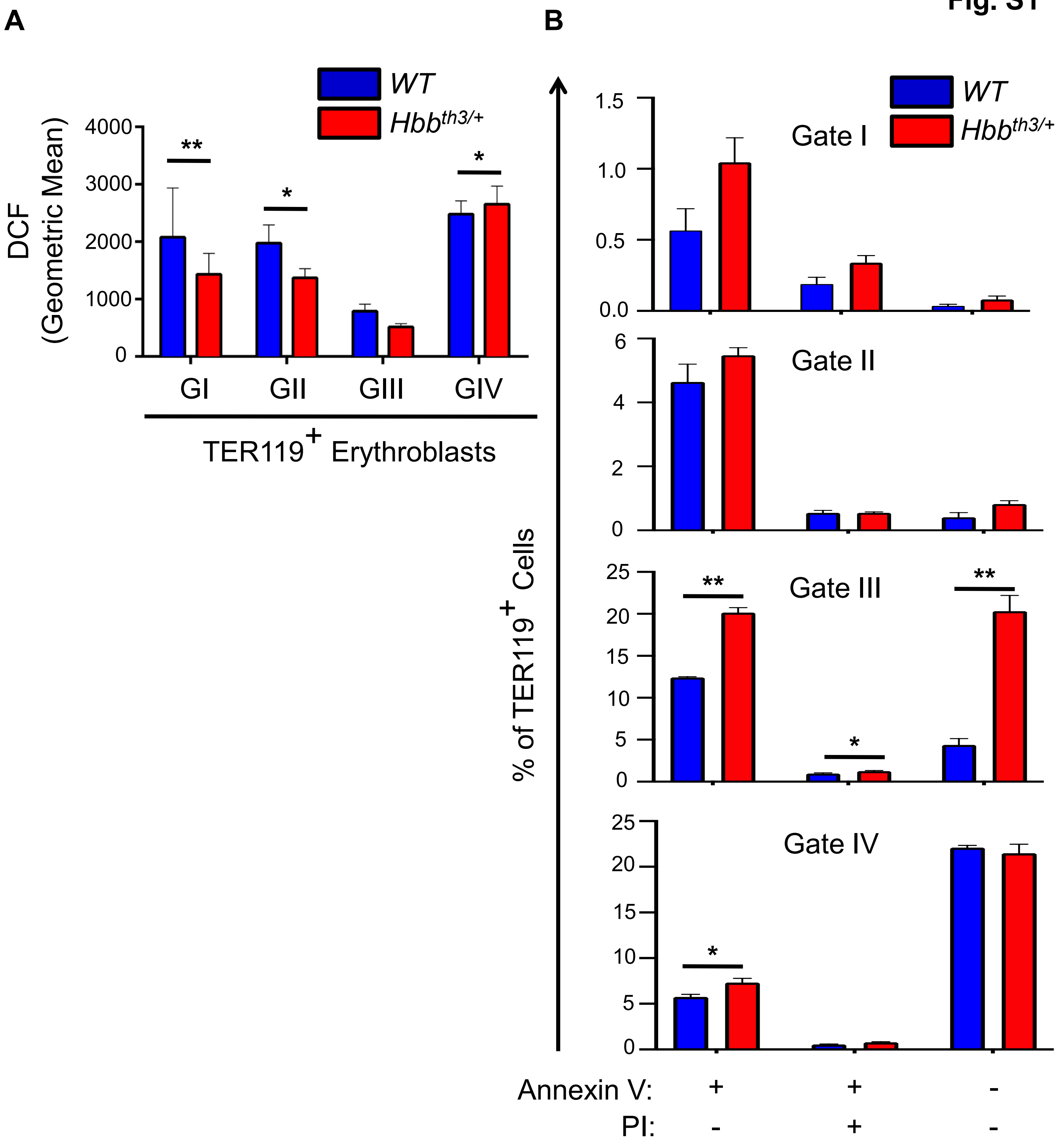
A
WT
Hbbth3/+
DCF
(Geometric Mean)
4000
3000
2000
1000
0
**
*
*
GI
GII
GIII
GIV
TER119+ Erythroblasts
B
1.5
1.0
0.5
0.0
Gate I
6
4
2
0
Gate II
25
20
15
10
5
0
Gate III
**
*
**
Gate IV
*
% of TER119+ Cells
Annexin V: + + -
PI: - + -

Fig. S2

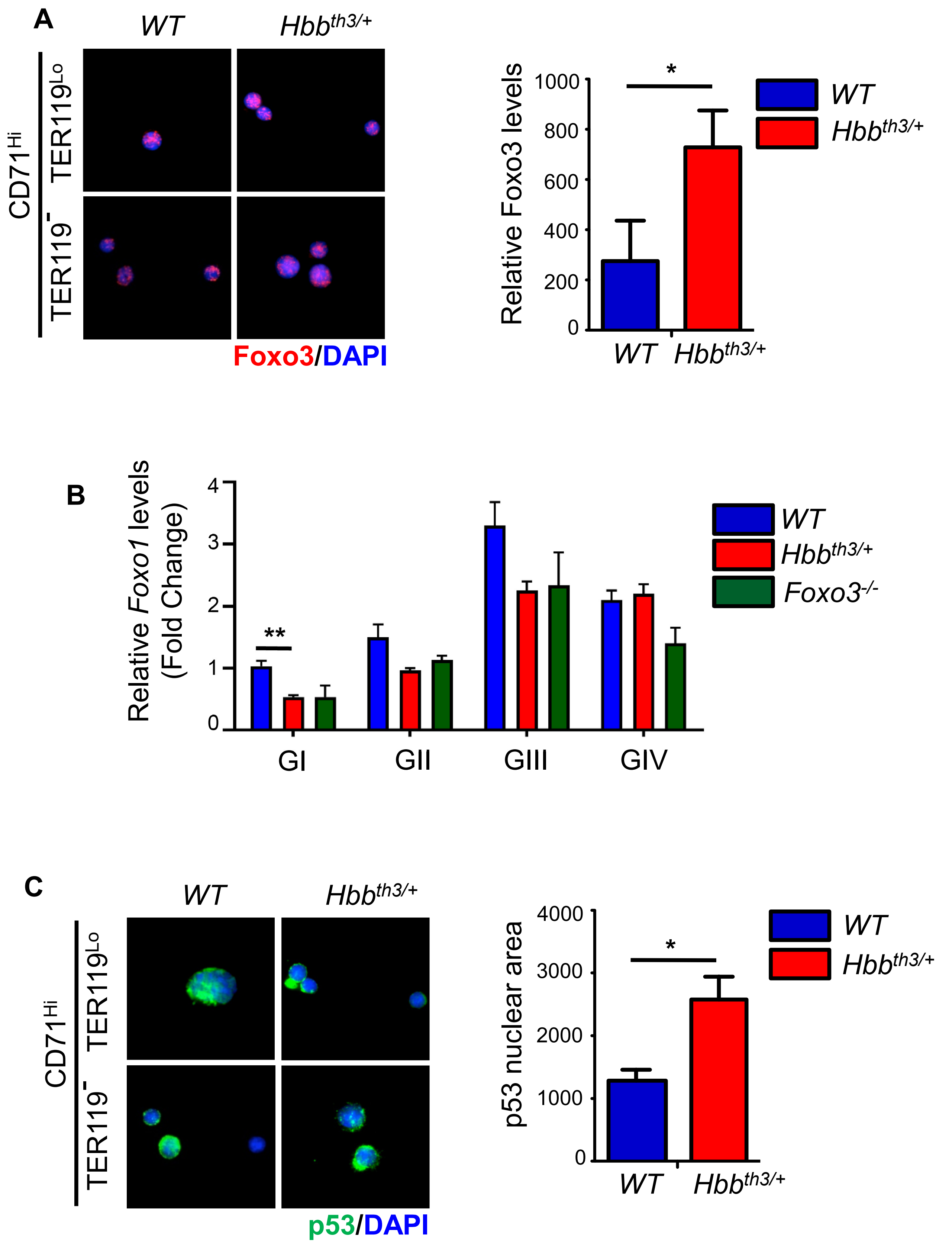
A
WT
Hbbth3/+
CD71Hi
TER119Lo
TER119-
Foxo3/DAPI
Relative Foxo3 levels
1000
800
600
400
200
0
*
WT
Hbbth3/+
B
Relative Foxo1 levels
(Fold Change)
4
3
2
1
0
**
GI
GII
GIII
GIV
WT
Hbbth3/+
Foxo3-/-
C
WT
Hbbth3/+
CD71Hi
TER119Lo
TER119-
p53/DAPI
p53 nuclear area
4000
3000
2000
1000
0
*
WT
Hbbth3/+

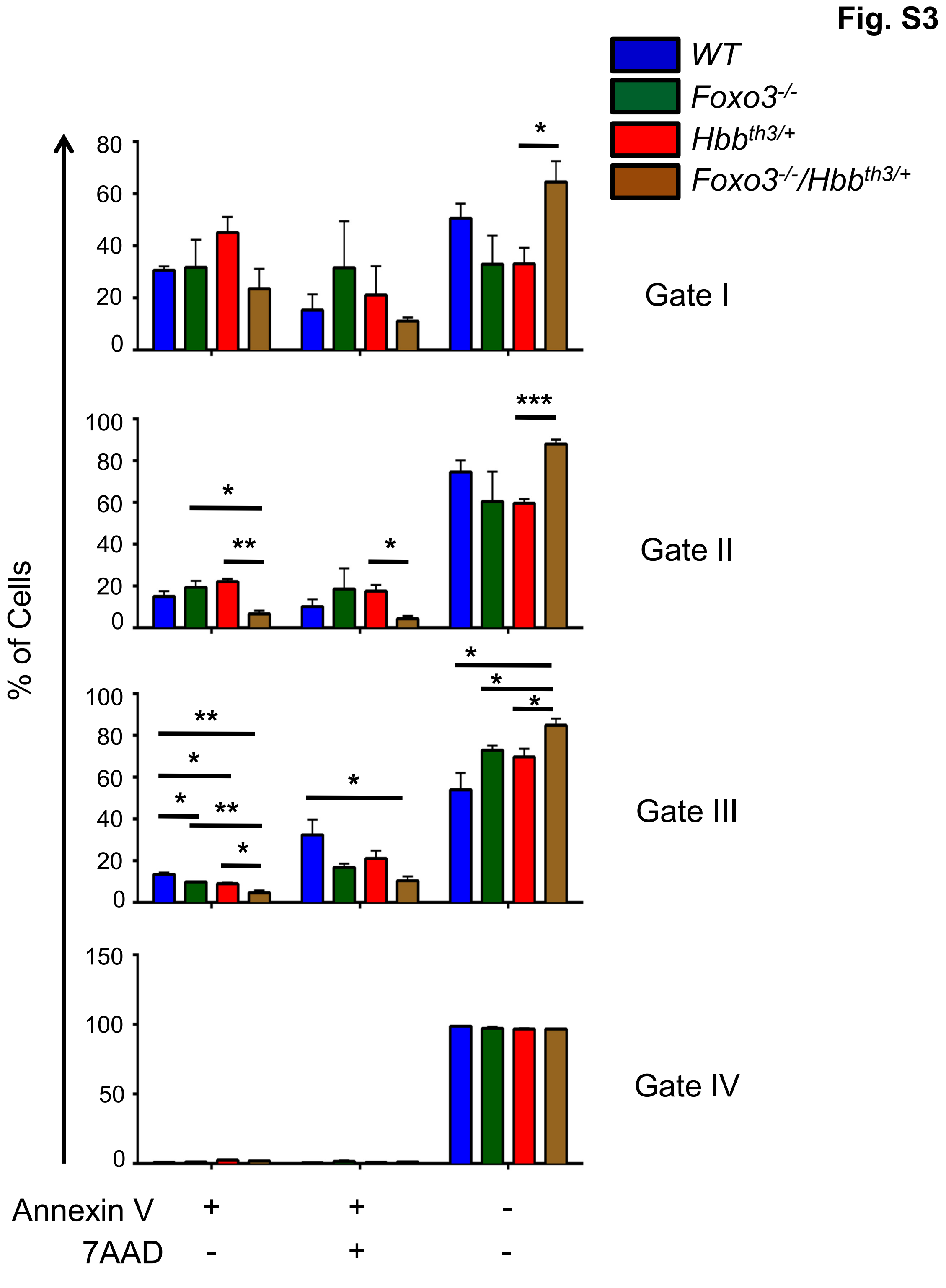
Fig. S3
WT
Foxo3-/-
Hbbth3/+
Foxo3-/-/Hbbth3/+
% of Cells
Gate I
Gate II
Gate III
Gate IV
Annexin V + + -
7AAD - + -
0
20
40
60
80
100
150
50

Fig. S4

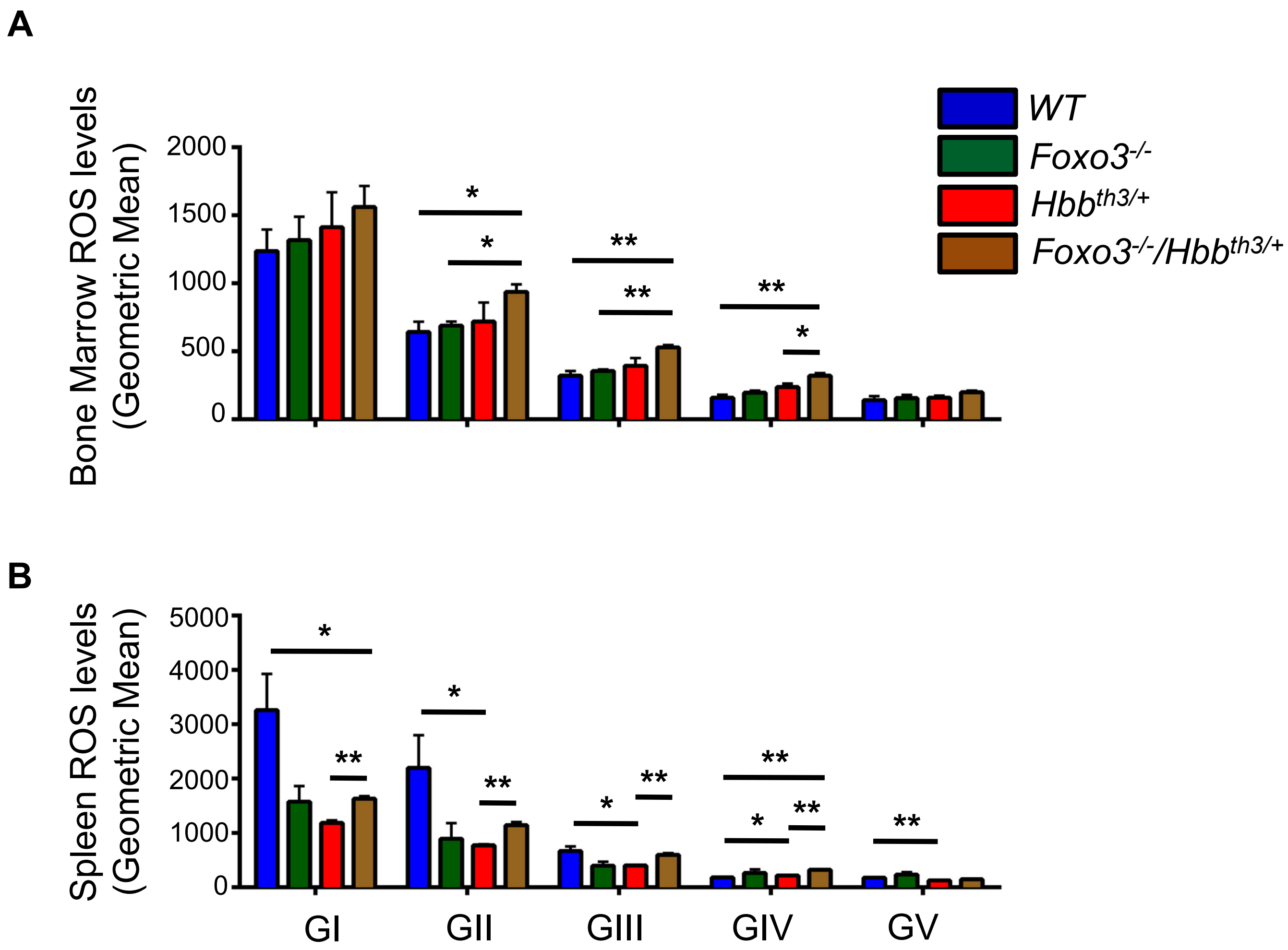
A
Bone Marrow ROS levels (Geometric Mean)
2000
1500
1000
500
0
WT
Foxo3-/-
Hbbth3/+
Foxo3-/-/Hbbth3/+
B
Spleen ROS levels (Geometric Mean)
5000
4000
3000
2000
1000
0
GI
GII
GIII
GIV
GV

Fig. S5

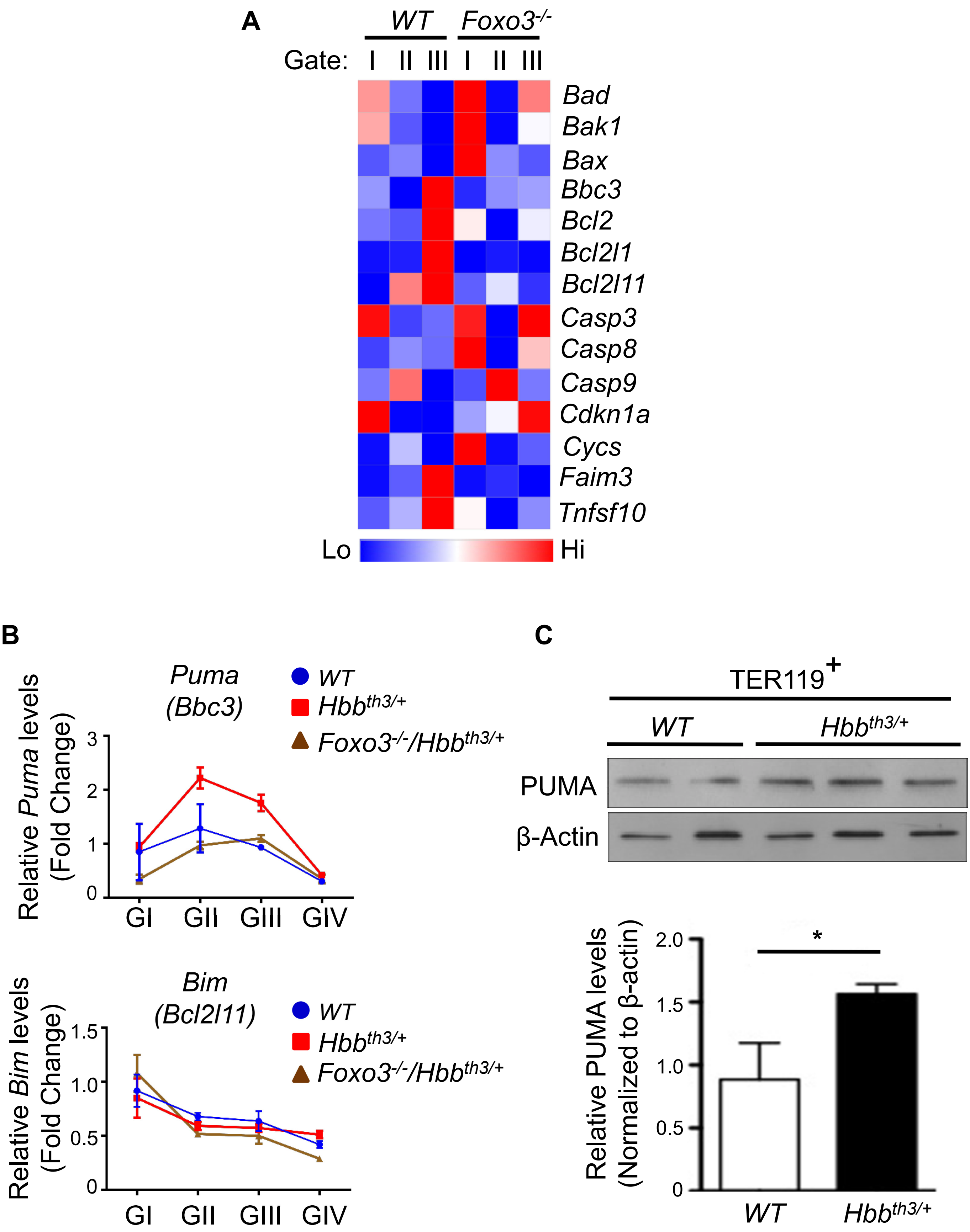
A
WT
Foxo3-/-
Gate: I II III I II III
Bad
Bak1
Bax
Bbc3
Bcl2
Bcl2l1
Bcl2l11
Casp3
Casp8
Casp9
Cdkn1a
Cycs
Faim3
Tnfsf10
Lo
Hi
B
Puma (Bbc3)
WT
Hbbth3/+
Foxo3-/-/Hbbth3/+
Relative Puma levels (Fold Change)
3
2
1
0
GI GII GIII GIV
Bim (Bcl2l11)
WT
Hbbth3/+
Foxo3-/-/Hbbth3/+
Relative Bim levels (Fold Change)
1.5
1.0
0.5
0
GI GII GIII GIV
C
TER119+
WT
Hbbth3/+
PUMA
β-Actin
Relative PUMA levels (Normalized to β-actin)
2.0
1.5
1.0
0.5
0
*
WT
Hbbth3/+

Fig. S6

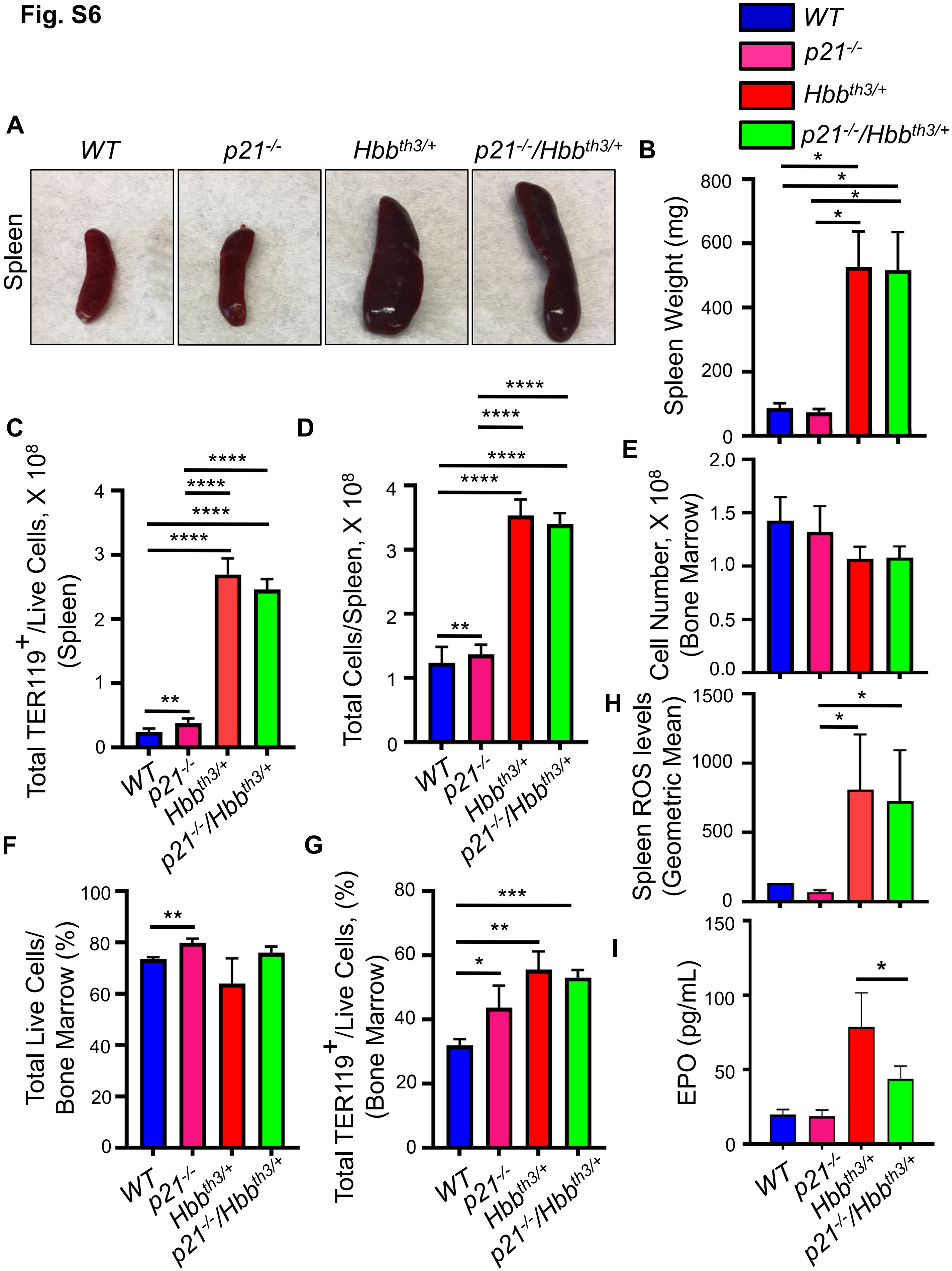
WT
p21-/-
Hbbth3/+
p21-/-/Hbbth3/+
A
Spleen
B
Spleen Weight (mg)
800
600
400
200
0
*
C
Total TER119+/Live Cells, X 10^8 (Spleen)
4
3
2
1
0
****
**
D
Total Cells/Spleen, X 10^8
E
Cell Number, X 10^8 (Bone Marrow)
2.0
1.5
1.0
0.5
0.0
F
Total Live Cells/ Bone Marrow (%)
100
80
60
40
20
G
Total TER119+/Live Cells, (%) (Bone Marrow)
***
H
Spleen ROS levels (Geometric Mean)
1500
1000
500
I
EPO (pg/mL)
150
100
50

Fig. S7

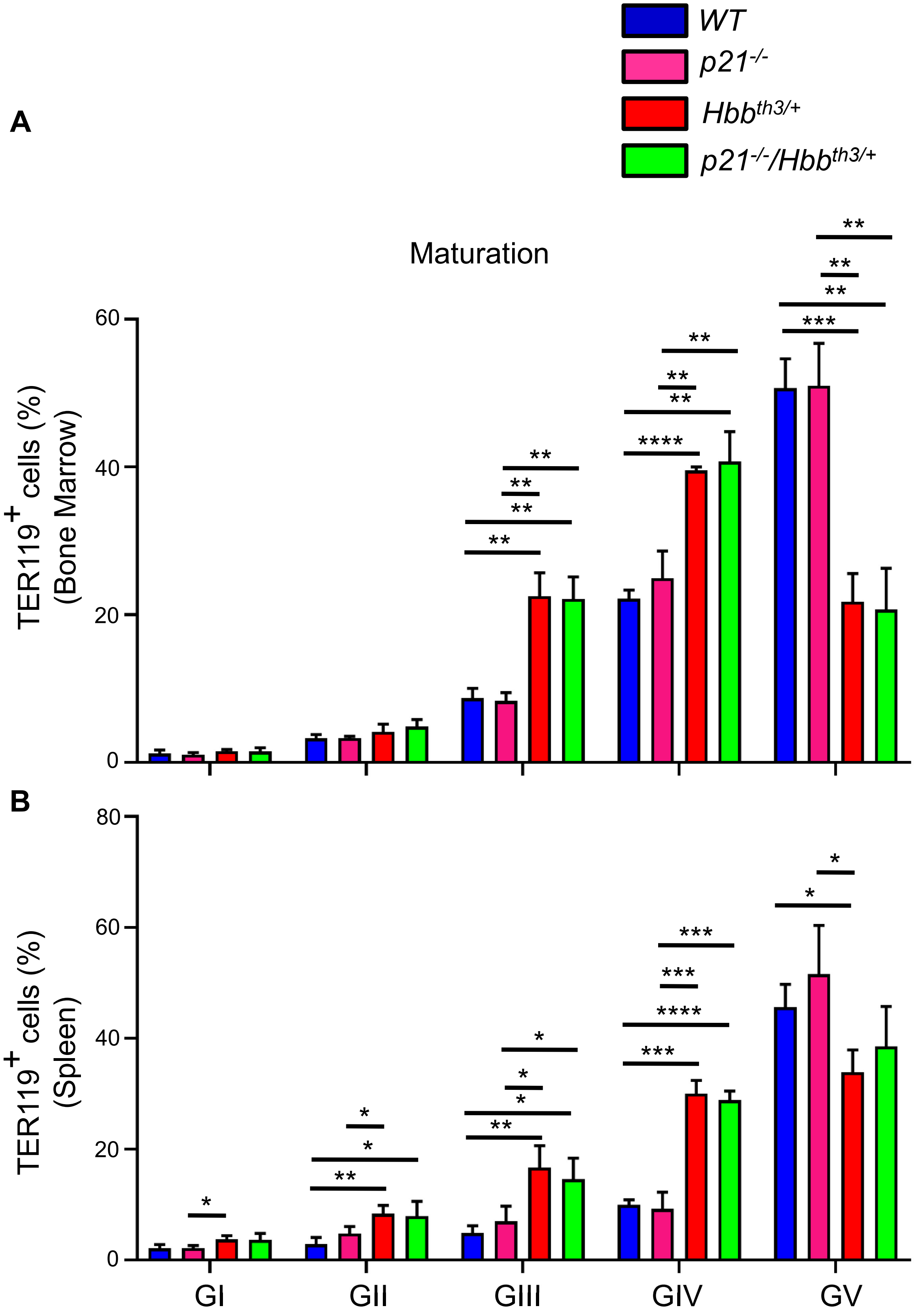
A
B
WT
p21-/-
Hbbth3/+
p21-/-/Hbbth3/+
Maturation
TER119+ cells (%) (Bone Marrow)
TER119+ cells (%) (Spleen)
0
20
40
60
80
GI
GII
GIII
GIV
GV

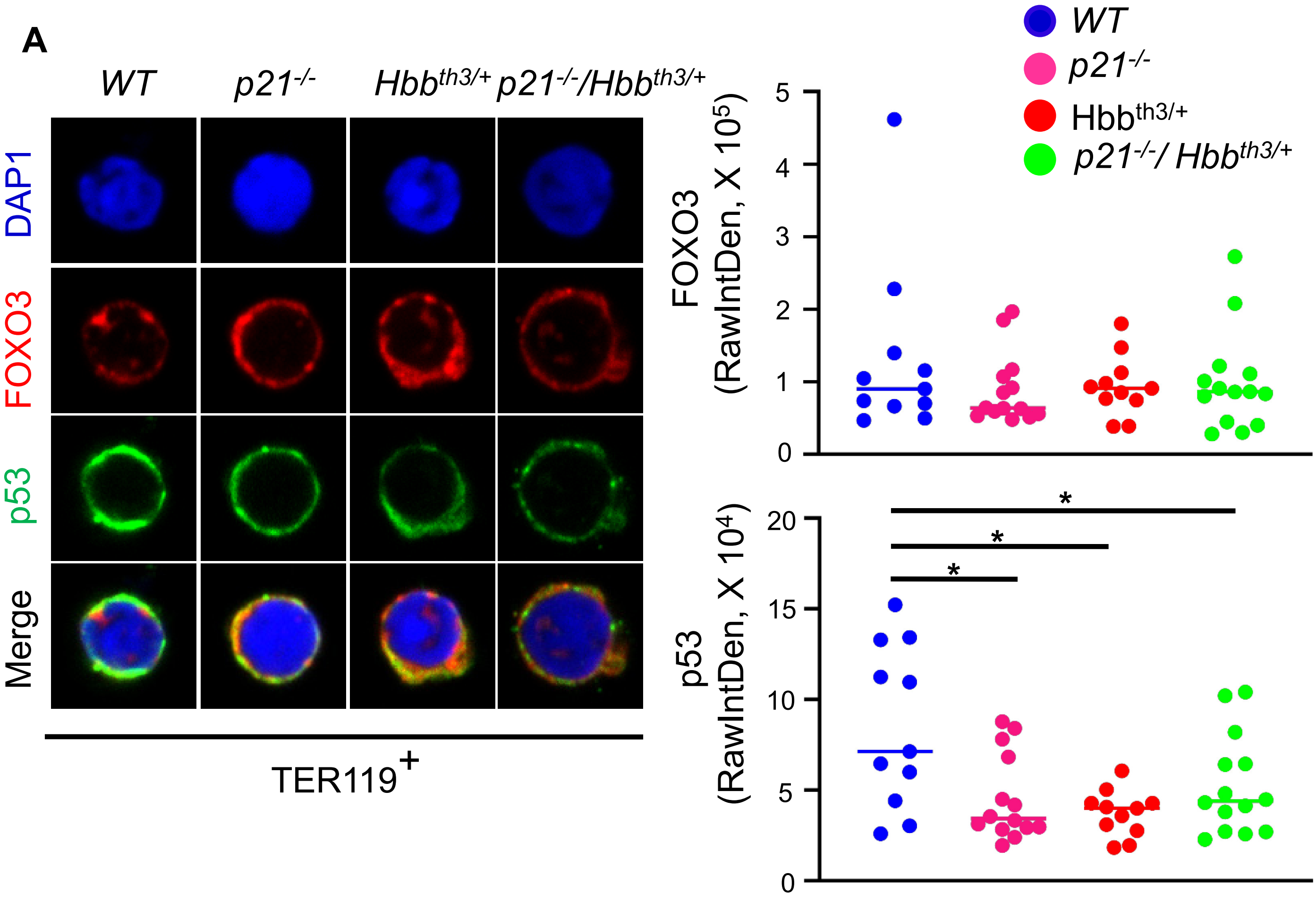
A
WT
p21-/-
Hbbth3/+
p21-/-/Hbbth3/+
DAP1
FOXO3
p53
Merge
TER119+
WT
p21-/-
Hbbth3/+
p21-/-/ Hbbth3/+
FOXO3 (RawIntDen, X 105)
p53 (RawIntDen, X 104)
*
*
*

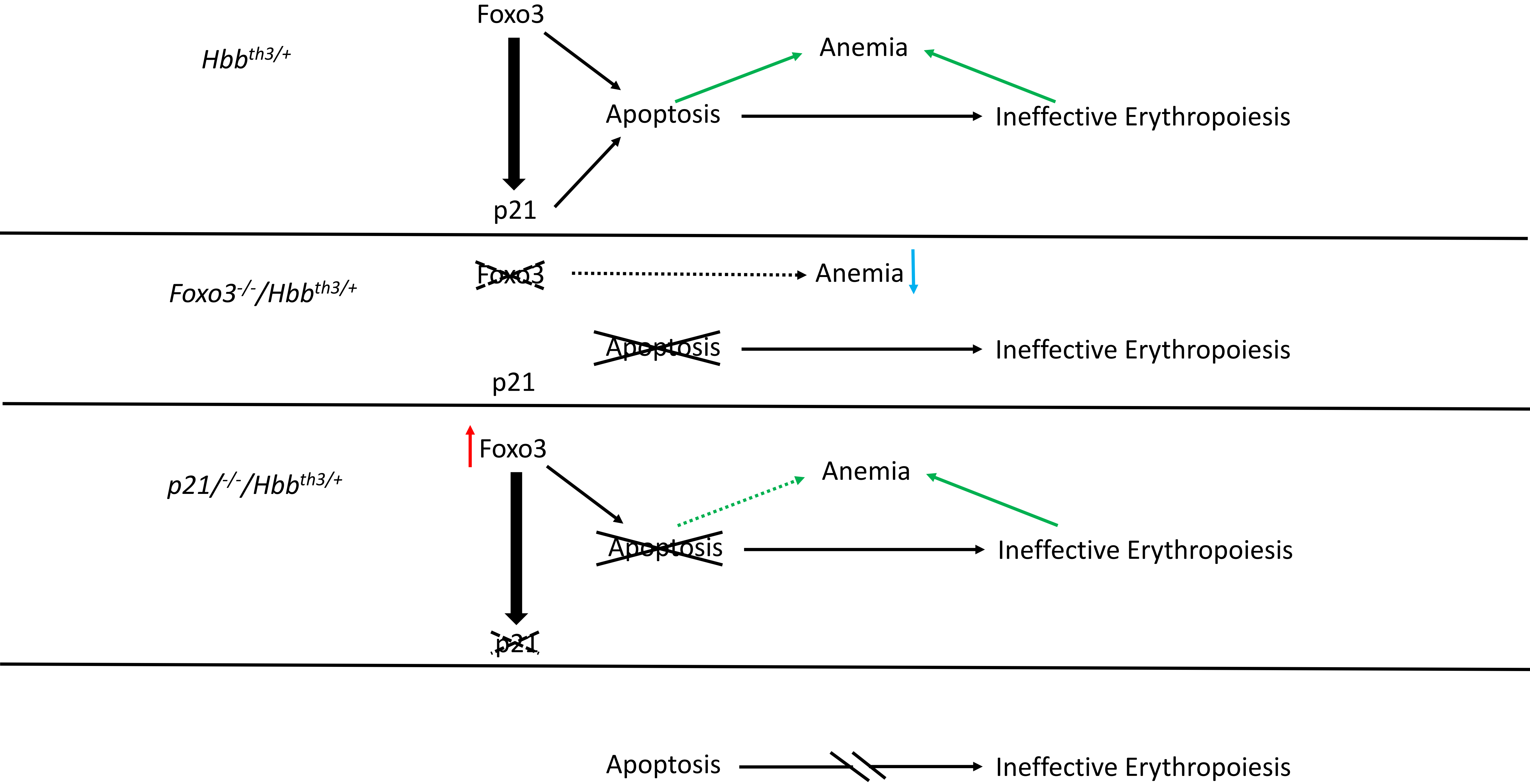

Hbb th3/+
Foxo3
p21
Apoptosis
Anemia
Ineffective Erythropoiesis
Foxo3-/-/Hbb th3/+
Foxo3
Anemia
Apoptosis
Ineffective Erythropoiesis
p21
p21/-/-/Hbb th3/+
Foxo3
Anemia
Apoptosis
Ineffective Erythropoiesis
p21
Apoptosis
Ineffective Erythropoiesis
